# Thioredoxin Reductase 1 inhibition triggers ferroptosis in KRAS-independent lung cancers

**DOI:** 10.1101/2025.07.25.666783

**Authors:** Cristina Andreani, Caterina Bartolacci, Margherita Melegari, Nicola Sargentoni, Lorenzo Luciani, Agnese Marucci, Roberta Galeazzi, Gina M. DeNicola, Jessica Kilgore, Noelle Williams, Stefano Berto, Massimiliano Gaetani, Prasad Pattabhi, Sagid S. Osman, Ahmad T. Mansour, Stefania Pucciarelli, Rossana Galassi, Pier Paolo Scaglioni

## Abstract

Lung cancers that harbor wild type KRAS (KRAS-WT) represent a molecularly diverse subset of tumors that often lack targeted therapeutic options. Using synthesized gold(I)-based inhibitors, a multi-omics approach, and functional validation, we identified Thioredoxin reductase 1 (TXNRD1), encoding as a selective vulnerability in KRAS-WT and oncogenic KRAS mutant (KM)-independent lung cancer (LC). Mechanistically, TRXR1 blockade induces ferroptosis through glutathione depletion, lipid reactive oxygen species (ROS) accumulation, and HMOX1-dependent iron overload in KRAS-WT LC both in vitro and in vivo. Furthermore, while KM LC cells are intrinsically resistant to TRXR1 inhibition, KMLC cells that acquire resistance to KRAS inhibitors (KRASi) undergo a redox shift that renders them sensitive to TRXR1 inhibition, uncovering a potential novel therapeutic vulnerability in KRASi-refractory tumors. These findings establish TRXR1 as a targetable redox checkpoint in KRAS-WT and KRASi-resistant lung cancers and support further development of TRXR1 inhibitors.

## Main

Lung cancer (LC) is a leading cause of cancer-related deaths^1^ and despite the great advancement made in the last decades, therapy resistance continues to be a challenge in treating this cancer^2^.

Ferroptosis, a programmed cell death characterized by iron- and reactive oxygen species (ROS)-dependent lipid oxidation^3^, has been implicated in several biological contexts, from physiology to disease^4–7^. Ferroptosis has gained attention as a potential target for cancer therapy as several studies have pointed out that cancer cells employ several strategies to suppress ferroptosis and withstand cancer treatments^8–12^.

Our group has previously described that LC cells harboring oncogenic *KRAS* mutations (KM) exploit fatty acid synthase (FASN) to feed oxidation resistant fatty acids into the Lands cycle and deflect ferroptosis^11^. Therefore, pharmacological inhibition of FASN induces ferroptosis in KMLC and can be used as a targetable vulnerability in KMLC patients (NCT03808558). Despite these promising results in KMLC, which accounts for about 30% of all non-small cell lung cancers (NSCLC), FASN inhibition (FASNi) was not effective in preclinical models of LC with KRAS-wild type (KRAS-WT), which represents the remaining 60-70% of NSCLC.

We demonstrated not only that KM expression in KRAS-WT was necessary and sufficient to establish the dependency on FASN, but also that KMLC cells that became resistant to KM ablation were also resistant to FASNi^11^. This observation suggested that LC cells may evolve different mechanisms to evade ferroptosis based on KRAS mutational status and/or dependency.

Cellular redox homeostasis is primarily regulated by the thioredoxin (TRX)- and glutathione (GSH)-dependent systems. The functions of these two systems are complementary and often synergize with or compensate for each other to maintain the redox balance in different physiological and disease contexts^13–18^. The TRX system, comprising TRXs (genes *TXNs*), thioredoxin reductases (TRXR1, genes *TXNRD*), and peroxiredoxins (PRDXs), plays a crucial role in antioxidant defense, DNA synthesis, and redox-regulated processes such as signal transduction, gene expression, and cell death^19,20^. The TRXR1 system in mammals consists of three distinct isoforms: *TXNRD1* (cytoplasmic), *TXNRD2* (mitochondrial), and *TXNRD3*, which is primarily expressed in the testis. These enzymes are all selenoproteins containing a selenocysteine (Sec) residue, which is essential for their catalytic activity. They function as homodimers and utilize NADPH and FAD as cofactors to reduce the disulfide bonds in TRXs and a variety of other endogenous and exogenous substrates, including the dipeptide cystine^16^. While loss of *TXNs* and *TXNRDs* is either embryonically lethal or causes infertility in mammals^21,22^, cytosolic TRX1 (gene *TXN1*) and TRXR1 are required for cell proliferation^22–24^. Accordingly, several studies have proposed TRXR1 as a potential vulnerability during malignant transformation and cancer progression^8,17,18,25^. However, TRXR1 seems to have a context-dependent function and its precise role in NSCLC and how it impacts tumorigenesis based on the presence or absence of specific oncogenic drivers like KM remains unclear. Using publicly available datasets, novel tool compounds and preclinical models, we identified TRXR1 as a vulnerability in KRAS-WT and KM-independent LC. While we demonstrated that low GSH predisposes KRAS-WT LC to this vulnerability, TXNRD1 inhibitors trigger cell death by acting as class IV ferroptosis inducing compounds (FINs), in which uncontrolled downstream heme oxygenase (*HMOX1*) induction leads to iron overload and ferroptotic cell death^26^. Of note, we found that a similar mechanism can be exploited to target KMLC persister cells resistant to KRAS inhibition (KRASi). This evidence not only confirms that *KRAS* and *TXNRD1* dependencies are mutually exclusive but also suggests that TRXR1 inhibitors might be beneficial in second line or combination therapy to resensitize KMLC to KRASi.

## Results

### TXNRD1 is a genetic vulnerability in cancer cells that are either KRAS-WT or KRAS-independent

Our previous work established that KRAS-WT LC cells were intrinsically resistant to ferroptosis induced by FASN inhibition^11^. This evidence suggested the hypothesis that, in the absence of KM, LC cells might rely on alternative pathways to prevent ferroptosis. To test this hypothesis, we interrogated the CRISPR loss-of-function data available in the DepMap database^27^, to look for ferroptosis-related vulnerabilities specific for KRAS-WT LC cells (Figure 1A). We included key genes involved in iron metabolism, cysteine and GSH synthesis and metabolism, the FSP1/Vitamin K axis, NADPH, the dihydrofolate reductase (DHFR) pathway and the TRX/PRDX pathway as they are all involved in ferroptosis regulation and execution^5,6,28^. Among the genes analyzed, *TXNRD1* was the most significant hit for KRAS-WT as compared to KM LC cells (Figure 1A, B). According to this database, *KRAS* and *TXNRD1* dependency is mutually exclusive in human NSCLC (Fig. 1C), and in other cancers (Fig. 1D). We validated these data *in vitro*, both in 2D and in 3D conditions, where siRNA- or shRNA-mediated knockdown of *TXNRD1* induced cell death in KRAS-WT and EGFR-mutant (EGFR-MUT) LC lines (Supplementary Fig. 1A-E). Of note, opposite to what we previously reported regarding FASN-dependency, where ectopic expression of KM in KRAS-WT line H522 was sufficient to render them sensitive to FASN inhibition^11^, here it rendered them resistant to *TXNRD1*-knockdown (Supplementary Fig. 1 C-E). Overall, these data suggest that *TXNRD1* is a potential targetable vulnerability in KRAS-WT and KRAS-independent cancers.

**Figure 1.**
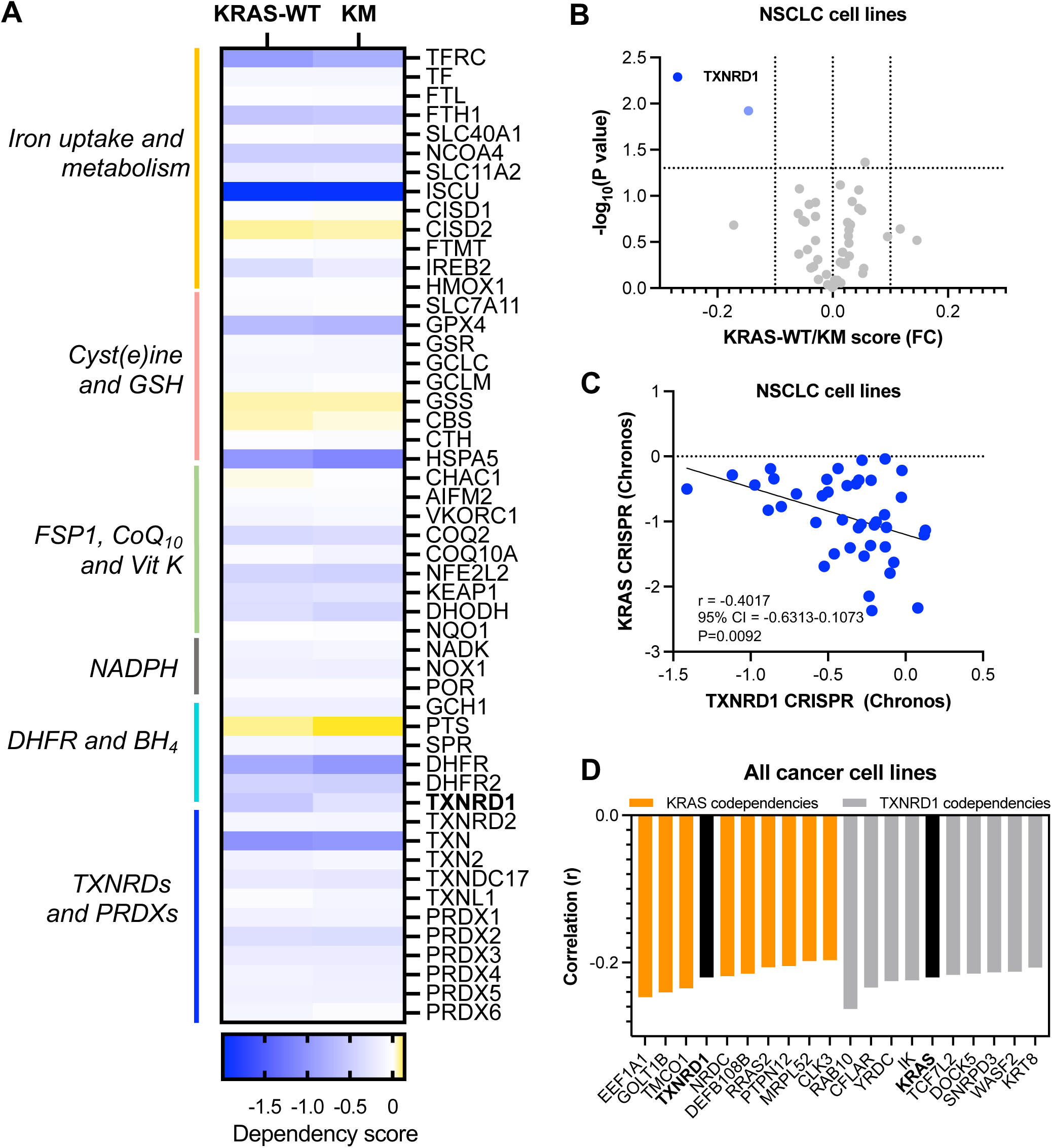
TXNRD1 is a genetic vulnerability in KRAS-WT and KRAS-independent cancer cells. A) Heatmap and volcano plot (B) representing the mean DepMap Dependency Scores (DepMap Public 23Q4+Score, Chronos) and their significance in KRAS-WT and KM lung cancer lines for the indicated genes. FC, fold change. P values represent multiple unpaired t test. C) Correlation plot between KRAS and TXNRD1 DepMap dependency in NSCLC cell lines. Correlation r, 95% CI and P values represent Pearson correlation. D) Top 10 KRAS and TXNRD1 DepMap co-dependencies in all cancer lines. Correlation r represents Pearson correlation.

### Newly synthesized gold(I) compounds are reversible TRXR1 inhibitors

To test whether pharmacological inhibition of TRXR1 can be exploited to target KRAS-WT LC cells, we used two in-house synthesized gold(I) inhibitors DM20 (4,5-dicyano-1H-imidazolate-1-yl)-(triphenylphosphane)-gold(I) and CS47 (4,5-dichloro-1H-imidazolate-1-yl)-(triphenylphosphane)-gold(I) (Fig. 2A). Our group described the TRXR1 inhibitory and anticancer activity of these compounds and similar analogs^29–32^, however their mechanism of action remained unknown. Here, employing *in silico* docking, we found that both compounds show reversible binding to the same pockets of human TRXR1 (Fig. 2B and Supplementary Fig. 2A). For both compounds the lowest energy cluster is positioned between FAD and NADPH sites (En = -8.03 kcal/mol for CS47; En = -7.59 kcal/mol for DM20). The binding is mediated by non-covalent interactions, including Leu385 and Ser386 (involved in OH–π interactions with the phenyl ring), and Phe228 (engaged in π–π stacking with the heterocyclic ring) (Fig. 2C). The second-best binding cluster (En = −7.42 kcal/mol for CS47; En = −7.33 kcal/mol for DM20) is in a shallower allosteric pocket positioned at a 180° rotation (Fig. 2B and Supplementary Fig. 2A). The interactions in this site are largely nonspecific, lacking ionic or hydrogen bonding. Being surface-exposed and not deeply embedded in the protein structure, this site likely supports weaker and less stable binding than cluster 1.

**Figure 2.**
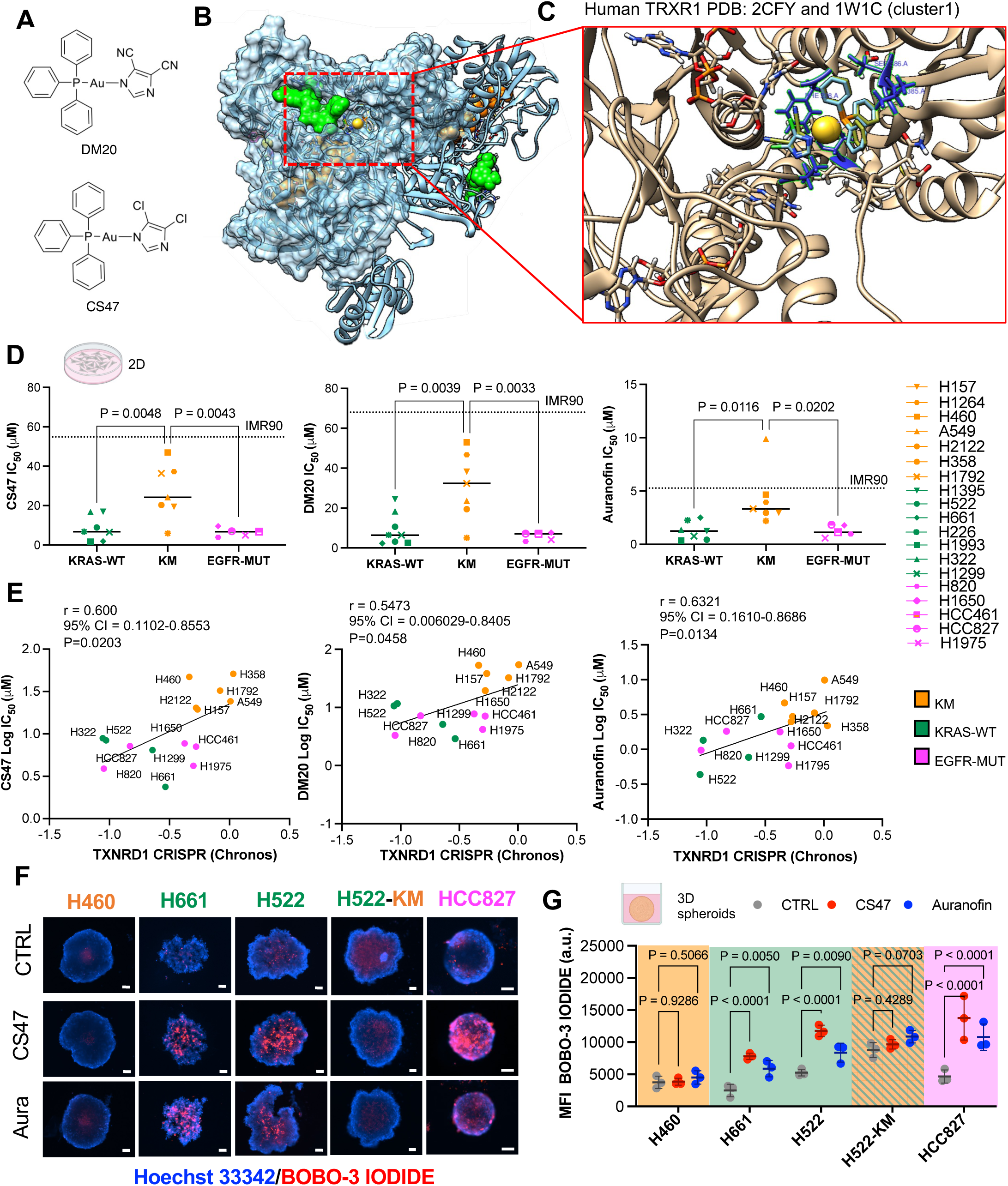
TRXR1 inhibition is selectively lethal to KRAS-WT lung cancer cells and spheroids. A) Chemical structure of the in-house synthesized gold(I) compounds DM20 and CS47 B) In silico docking of CS47 and DM20 cluster 1 with the human TRXR1 (PDB. 2CFY and 1W1C). The protein is represented in dimeric quaternary structure; a molecular surface is represented to better highlight the binding cleft. NADPH and FAD are represented as green and orange VdW spheres. Gold as VdW yellow sphere. C) Docking closeup showing the closest and interacting residues in blue. Namely: Leu385, Ser386 (OH—ρε with phenyl), Phe 228 ρε stacking with heterocyclic ring. NADPH and FAD are also represented as tubes, TRXR1 in ribbons, gold atoms as VdW yellow spheres. Hydrogen in the ligands are not shown for purpose of clarity. D) Plots representing the MTT IC_50_ of the indicated 2D cell lines treated with CS47, DM20 or auranofin for 48 hours. P values represent one-way ANOVA followed by Tukey’s multiple comparisons (n=2 independent experiments). E) Pearson correlation between the MTT IC_50_ of the indicated compounds and the DepMap genetic dependency on TXNRD1 (DepMap Public 23Q4+Score, Chronos). F) Live/Dead assay representative images and (G) quantification of images of 3D LC spheroids, treated as indicated for 48 hours. Scale bar, 100 µm. P values represent two-way ANOVA followed by Sidak’s multiple comparisons (n=3 independent replicates).

Although Cellular Thermal Shift Assay (CETSA)^33^ and Proteome Integral Solubility Assay (PISA)^34^ confirmed TRXR1 stabilization and TXNL1 degradation, consistent with effects previously reported for other TRXR1 inhibitors^35–39^ (Supplementary Fig. 2B–E), our docking analyses reveal key differences in the binding mechanisms of our gold(I) compounds compared to covalent inhibitors such as auranofin and TRi-1/TRi-2, which covalently modify the selenocysteine (Sec) and cysteine residues in the catalytic site or induce the formation of SecTRAPs^36–38^. Notably, the Cluster 1 binding site features a unique amino acid composition that differs from other homologous proteins^40^, such as glutathione reductase (GSR), suggesting that our compounds may offer greater specificity for TRXR1 than currently available covalent inhibitors. Before proceeding with the biological assays, we used ^1^H NMR spectroscopy (Supplementary Fig. 3) and Electrospray Ionization-Mass Spectrometry (ESI-MS) (Supplementary Fig. 4) to determine the overall stability of CS47 and DM20 and the identification of their likely species in solution. Analysis of the ^1^H NMR spectra of CS47 reveals that the peaks attributable to the aromatic rings of the PPh_3_ moiety as well as the ones of the imidazolyl ring (around 7.50 ppm) remain unchanged over a 72-hour period (Supplementary Fig. 3A). In contrast, DM20 displays variations of the chemical shifts of about 0.04 ppm (Supplementary Fig. 3B). Therefore, this analysis suggests that CS47 displays better stability than DM20. ESI-MS detected ionic adducts for both CS47 and DM20 (Supplementary Fig. 4). {[(PPh_3_)_2_Au] [Cl_2_ImAu]}^+^ was detected for CS47 at 1053 m/z (Supplementary Fig. 4A) while the ionic adduct {[(PPh_3_)_2_Au] [(CN)_2_ImAu]}^+^ was detected at 1035 m/z for DM20 (Supplementary Fig. 4B). Upon incubation in buffered water/methanol solution for 24 hours, the ESI spectrum of CS47 displays the most intense peak at 475 m/z, this latter due to the displacement of the imidazolyl ligand from the complex by the ammonia derived from the buffer, [(NH_3_)AuPPh_3_]^+^ (Supplementary Fig. 4C), while, interestingly, in DM20 the full hydrolysis of the cyanide groups is detected by ESI MS, in which the species at 614.7 m/z, attributed to [(COOH)_2_ImAuPPh_3_]^+^, is the predominant peak even in the freshly prepared buffered solution (Supplementary Fig. 4D). Therefore, it is reasonable to conclude that the lower stability of DM20 observed in the ^1^H NMR spectra, can be attributed to the hydrolysis of the cyanide groups of the imidazolyl ligand in polar aqueous medium, affecting its overall stability, hence its activity. Given these results showing a better stability of CS47 in solution, we prioritized a more in depth *in vitro* and *in vivo* characterization of CS47.

### Pharmacological inhibition of TRXR1 is lethal to KRAS-WT lung cancer cells

We then assessed the activity of DM20, CS47 and auranofin (used as positive control for TRXR1 inhibition) in a panel of 20 human cell lines (LC and normal lung fibroblasts IMR90) in 2D conditions (Fig. 2D). Despite two independent assays confirmed that TRXR1 activity was inhibited in both groups to a similar extent (Supplementary Fig. 5A, B), KRAS-WT LC cells were highly sensitive to TRXR1 inhibition regardless of the drug used, while KMLC cell lines exhibit resistance (Fig. 2D and Supplementary Fig. 5C). A similar outcome was seen in redox immunoblots showing that two TRXR1 targets, TRX1 and PRDX1, were oxidized upon TRXR1 inhibition only in the sensitive KRAS-WT LC cells (Supplementary Fig. 5D). Noteworthy, the resistant behavior is specific for KMLC cells only, as cell lines driven by other oncogenes such as EGFR-mutant (EGFR-MUT) were also sensitive to the inhibition of TRXR1 (Fig. 2D and Supplementary Fig. 5C). These results reinforce the notion that TRXR1 is dispensable in KMLC as these tumor cells rely on alternative KM-driven programs to maintain their redox homeostasis, as we previously described^11^. Notably, although auranofin exhibits lower overall IC_50_ values (1.27 µM in KRAS-WT; 1.26 µM in EGFR-MUT; 4.27 µM in KM) compared to our reversible gold(I) compounds CS47 (6.74 µM in KRAS-WT; 6.78 µM in EGFR-MUT; 24.22 µM in KM) and DM20 (6.46 µM in KRAS-WT; 7.09 µM in EGFR-MUT; 31.21 µM in KM)—a difference expected due to auranofin’s covalent and irreversible mechanism—our compounds demonstrate superior selectivity. Specifically, CS47 and DM20 more effectively discriminate between sensitive (KRAS-WT and EGFR-MUT) and resistant KM LC cells and show substantially reduced cytotoxicity toward non-tumor lung fibroblasts (IMR90: CS47 IC_50_= 54.85 µM; DM20 IC_50_ = 64.02 µM; auranofin IC_50_ = 5.27 µM). These findings suggest that CS47 and DM20 exhibit enhanced TRXR1 specificity and reduced off-target effects compared to auranofin.

In contrast with what other studies have reported, but consistent with the DepMap data summarized in Fig.1, we found a positive correlation between sensitivity to the pharmacological inhibition of TRXR1 and the genetic dependency on *TXNRD1*, suggesting that inhibition or loss of this pathway are equally detrimental or tolerated in LC lines (Fig. 2E). As 2D and 3D culture conditions can alter sensitivity to treatments, ferroptosis and ROS^41,42^, we tested the effect of TRXR1 inhibition on LC spheroids (Fig. 2F, 2G). 3D cultures recapitulated the results obtained in 2D conditions, with KRAS-WT and EGFR-MUT LC spheroids exhibiting the highest degree of cell death, while KMLC cells were refractory to the treatment (Fig. 2F, 2G). As shown for *TXNRD1* knock-down (Supplementary Fig. 1), we demonstrated that KM overexpression was also sufficient to induce resistance to TRXR1 inhibition in KRAS-WT LC spheroids (Fig. 2F, 2G). Overall, these results indicate that TRXR1 inhibition can be exploited to kill LC cells in absence of KM.

### TRXR1 inhibition induces ferroptosis in KRAS-WT LC cells *in vitro*

To test whether ferroptosis is the mechanism underlying the cytotoxic effects exerted by TRXR1 inhibition, we first determined whether CS47, DM20 or auranofin cause the oxidation of the lipid probe C11-BODIPY, as an indirect measurement of lipid peroxidation in KRAS-WT and KM LC lines (Fig. 3A, 3B). While KMLC cells display either unchanged or decreased C11-BODIPY oxidation, suggesting again that they are not relying on TRXR1 to maintain redox homeostasis, DM20, CS47 and auranofin induced a significant C11-BODIPY oxidation in KRAS-WT LC lines (Fig. 3A, 3B). Next, we used untargeted liquid chromatography and tandem mass spectrometry (LC-MS/MS) to assess the endogenous lipidomic changes in H522 KRAS-WT LC cells. Consistent with the C11-BODIPY data, both DM20 and CS47 induced enrichment of phospholipids (PL) containing polyunsaturated fatty acids (PUFA), mainly phosphatidylcholines (PC) and phosphatidylethanolamines (PE), which are the preferred substrates of lipid peroxidation during ferroptosis^43,44^ (Fig. 3C-3E). As previously reported for other ferroptosis inducers^11,45,46^, DM20 and CS47 also induced accumulation of lysophosphatidylcholines (LPC), mainly containing saturated and monounsaturated acyl chains (SFA and MUFA), which result from the cleavage of oxidized PUFA-PC during ferroptosis. We also observed an enrichment of PUFA within triacylglycerols (TG) (Fig. 3C-3E). This is likely a mechanism to sequester excess PUFA into TG stored in lipid droplets to limit their incorporation into PL, the main substrate of lipid peroxidation^47^. Consistent with KRAS-WT LC cells undergoing ferroptosis, we not only detected a significant increase in intracellular malondialdehyde (MDA) and ferrous iron (Fe^2+^) (Supplementary Fig. 6A, 6B), but we also demonstrated that the ferroptosis inhibitor ferrostatin-1 (Fer-1) and the iron chelator deferoxamine (DFO) rescued the cell death induced by auranofin or our lead compound CS47 (Fig. 3F and Supplementary Fig. 6C). As expected, a similar rescuing effect was achieved by L-cysteine and N-acetyl cysteine (NAC) (Supplementary Fig. 6D), that can either act directly as antioxidant by using the free thiol (-SH) groups or indirectly by providing cysteine for GSH production bypassing the need for TRXR1 function.

**Figure 3.**
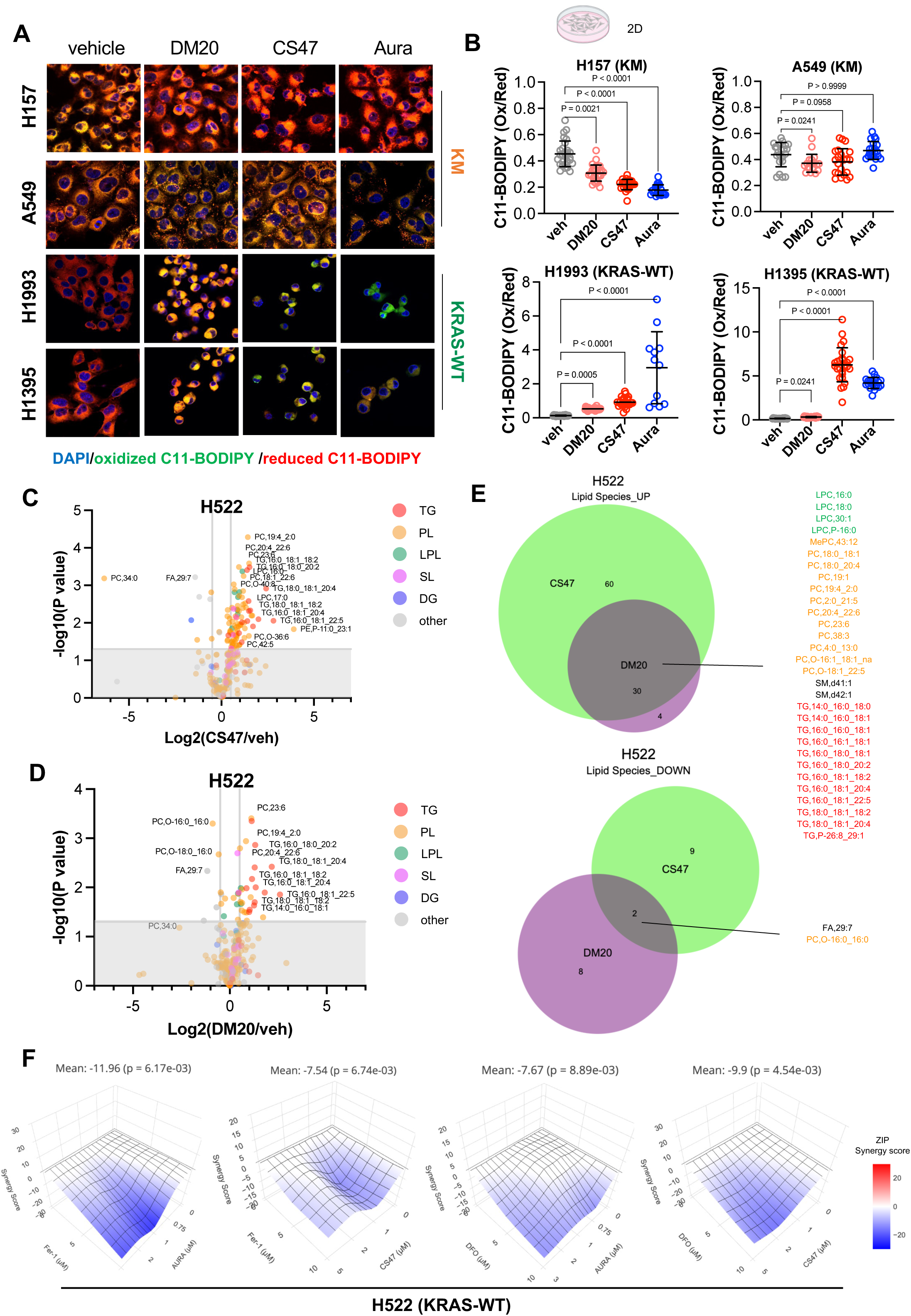
TRXR1 inhibition induces ferroptosis in KRAS-WT lung cancer cells. A) Representative images and quantification (B) of C11-BODIPY staining in the indicated lung cancer cell lines treated as indicated for 24 hours. P values represent one-way ANOVA followed by Dunnett’s multiple comparisons. C) Volcano plot of the differentially abundant lipid species detected via LC-MS lipidomics in H522 cells treated with 2 µM CS47 or DM20 (D) over vehicle for 24 hours. E) Venn diagrams showing the overlap of significant lipid species (P<0.05) significantly enriched or depleted in H522 cells treated with 2 µM CS47 or DM20 (n=4 independent replicates). F) Synergy plots of the indicated treatment combinations in H522 cells assessed as MTT assay at 48 hours (n=2 independent experiments/each combination). P values for the average synergy scores were derived by bootstrapping of the dose–response matrix in www.synergyfinderplus.org.

### Noncovalent TRXR1 inhibition is well tolerated *in vivo* and induces ferroptosis in KRAS-WT LC tumors

Before testing our lead compound CS47 in a tumor model we established its pharmacokinetic (PK) profile in NCG mice. To this end, we adapted a previously described diethyldithiocarbamate (DDTC) derivatization method^48^, converting plasma CS47 into the CS47Au(DDTC)_2_ adduct, which was subsequently detected by LC-MS (Methods, Supplementary Fig. 7A-C). After i.p. administration of 10 mg/kg, CS47 reached a maximum concentration (C_max_) of 20.33 μg/ml at 90 minutes (Supplementary Fig. 7B-C). CS47 plasma concentration decreased to a concentration of 14.26 μg/ml at 24 hours, but remained stable therafter, with a half-life (T_1/2_) of approximately 2.6 days (3803 min) (Supplementary Fig. 7B-C).

Then, we performed an acute toxicity analysis, in which mice were challenged with a single dose of 3, 5 and 10 mg/kg CS47 or vehicle, and then monitored for 3 weeks post administration. Even though normal activity and survival were noted for the 3 and 5 mg/kg doses at 24 hours, animals administered CS47 at 10 mg/kg showed slightly reduced activity immediately after compound administration and were found deceased in their cages 25 days after compound delivery. Therefore, we determined that 5 mg/kg was the optimal maximum tolerated dose (MTD) that did not produce any death, weight loss or changes in activity within the 25 days post treatment (Supplementary Table 1). Using these parameters, we established that 3 mg/kg i.p. every 3 days as the proper treatment regimen for in vivo anti-tumor studies to be within the optimal plasma range and MTD of CS47 (Supplementary Fig. 7A-C, Supplementary Table 1). Briefly, NCG mice bearing H522 xenografts received either CS47 or auranofin with or without the ferroptosis inhibitor, Liproxstatin-1 (Lip-1) as previously described^11,49^, over a period of 25 days. As reported in Fig. 4A, CS47 and auranofin produced a significant anti-tumor effect in H522 xenografts, while Lip-1 completely rescued the tumor growth in both treatment groups, confirming that TRXR1 inhibition induces ferroptosis in KRAS-WT LC tumors *in vivo*.

**Figure 4.**
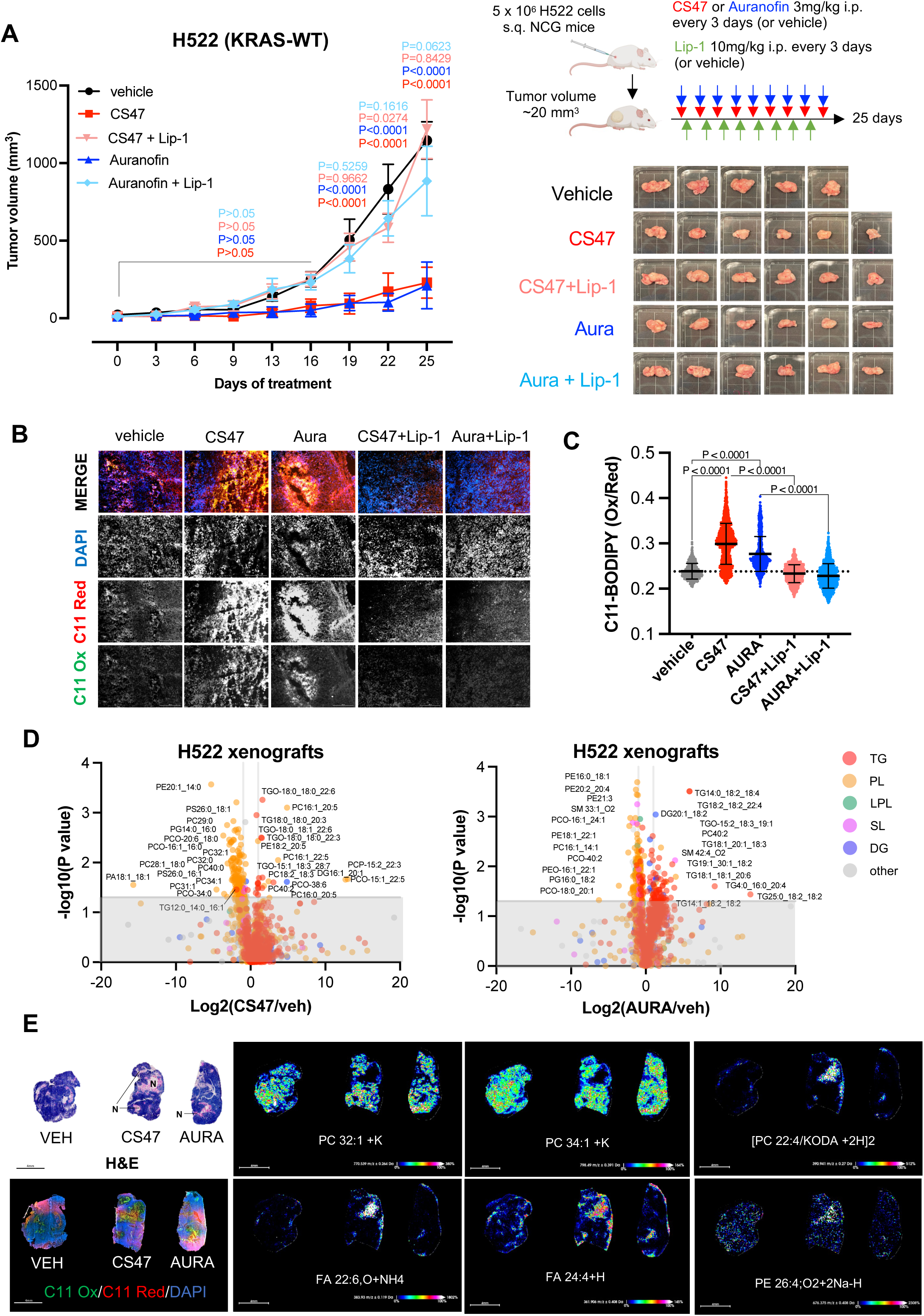
TRXR1 inhibition induces ferroptosis in KRAS-WT lung cancer in vivo. A) Tumor growth curves of H522 xenografts (n=5-6 mice/group) treated as indicated. P values represent two-way ANOVA followed by Dunnett’s multiple comparisons. Treatment scheme and post dissection images of H522 xenografts are reported on the right. B) C11-BODIPY staining of H522 xenografts treated as indicated. C) Plots represent quantification of C11-BODIPY staining H522 cells pooled from xenografts; n= 4 mouse per group. P values were calculated using one-way ANOVA followed by Sidak’s tests. D) Volcano plots showing the differentially represented lipid species identified by LC-MS lipidomics in H522 xenografts (n= 3 mice for vehicle and 4 mice for CS47 or auranofin). P values were calculated using multiple two-tailed t tests followed by Benjamini, Krieger and Yekutieli FDR. E) Representative images of matched H522 xenografts treated as indicated and stained with H&E, C11-BODIPY or assessed by MALDI-IMS on H522 xenografts treated as indicated. Rainbow scale represents % ion intensity normalized against the total ion count (TIC). Observed m/z and mass error (ppm) values are indicated for each lipid species. N indicates the necrotic areas.

These results were corroborated by C11-BODIPY staining, showing increased oxidation of the lipid probe in the CS47 and auranofin groups, while co-treatment with Lip-1 rescued C11-BODIPY oxidation *in vivo* (Fig. 4B, C). Also, consistent with our findings *in vitro* (Fig. 3), LC/MS-MS lipidomic analysis showed accumulation of PUFA-PL and PUFA-TG also in tumors treated with CS47 and auranofin (Fig. 4D). Moreover, employing MALDI-IMS as we previously described^11^, we identified that those area which appeared to be majorly affected by necrosis and C11-BODIPY oxidation, show a pro-ferroptosis lipid profile, with low MUFA-PC (e.g. PC 32:1, PC 34:1), but enrichment in PUFA-PL (e.g. PE 26:4), oxidatively truncated PUFA-PL^50^ (e.g. PC 22:4/KODA) and free PUFA such as docosahexaenoic acid (DHA, FA 22:6) and adrenic acid (FA 22:4) (Fig. 4E)^44,51^.

Although CS47 and auranofin exhibited comparable anti-tumor efficacy without causing significant weight loss in the animals (Supplementary Fig. 7D), CS47 demonstrated a more favorable hematological toxicity profile (Supplementary Fig. 7E, F). In contrast, auranofin, alone or in combination with Lip-1, induced marked leukopenia and neutropenia by the end of the experiment (Supplementary Fig. 7E) while no significant differences in platelet or erythrocyte counts were observed among the treatment groups (Supplementary Fig. 7F). At the tissue level, no overt pathological alterations were observed in the kidneys, liver, or lungs, except for a tendency toward alveolar hemorrhage in the group treated with the combination of auranofin and Lip-1 (Supplementary Fig. 7G, H). Similar anti-tumor efficacy and toxicity profiles were observed in a more aggressive xenograft model using the EGFR-MUT human lung cancer cell line H1650^52^ (Supplementary Fig. 8), further supporting that CS47 is better tolerated than auranofin, despite eliciting comparable anti-tumor effects.

### TRXR1 inhibition activates stress-responsive GSH and iron regulatory programs in KRAS-WT LC

To gain deeper mechanistic insight into how TRXR1 inhibition triggers ferroptosis in sensitive cells, we performed bulk RNA-seq on four KM (A549, H460, H157, H1792), four KRAS-WT (H522, H661, H1993, H1299), and three EGFR-MUT (PC-9, HCC827, H1650) lung cancer cell lines treated with CS47, auranofin, or vehicle (Fig. 5 and Supplementary Fig. 9). As expected, the KRAS-WT and EGFR-MUT cells, identified as the most sensitive, exhibited the most pronounced transcriptomic changes, whereas the treatment-resistant KM LC cohort showed only a limited number of differentially expressed genes (DEGs) upon TRXR1 inhibition (Fig. 5A). Notably, common downregulated DEGs in the KRAS-WT and EGFR-MUT groups included genes involved in fatty acid and cholesterol biosynthesis (e.g., *FASN, SCD,* and *ELOVL6*) as well as selenoproteins and selenoprotein synthesis (*SELENOT, SELENON, SECISBP2*) downstream of TRXR1^6^ (Fig. 5A, B, Supplementary Fig. 9A). These changes align with the lipidomic, and oxidative stress profiles induced by CS47 and auranofin (Figs. 3 and 4) and support the concept that TRXR1 participates in selenoprotein synthesis producing selenide from both organic and inorganic sources^6^. Among the upregulated DEGs, there are ferroptosis-related genes involved in antioxidant response (including *TXNRD1, TXN, SRXN1, SOD1, GSR, NQO1)*, iron transport and heme metabolism (*FTL, SLC25A38, ALAS1, FECH1, BACH1, HMOX1)*, cysteine and GSH homeostasis (*SLC7A11, SLC3A2, GCLM, GCLC)* (Fig. 5A, B, Supplementary Fig. 9A). Most of these genes were validated at protein level via proteomics in H522 cells, suggesting that these changes lead to altered target protein levels (Supplementary Fig. 9B, C). This transcriptional reprogramming is reflected by the abundance of the metabolites involved in the synthesis of GSH and uncovers differences between KRAS-WT and KM LC cells treated with TRXR1 inhibitors (Supplementary Fig. 10). Indeed, while cystine and glycine levels decreased in both groups after 24 hours of treatment—consistent with their utilization for GSH synthesis and the maintenance of relatively stable cysteine levels—serine levels were uniquely upregulated in KM cells (Supplementary Fig. 10A). This finding suggests that, under TRXR1 inhibition, KM cells can rely on enhanced serine synthesis or uptake to support survival by supplying key precursors for nucleotide, lipid, and GSH biosynthesis, as previously reported for KM cancers^53–55^. This may explain why GSH levels at baseline were almost double in KM and H522-KM cells as compared to KRAS-WT cells, while KRAS-WT cells showed the most significant increase in GSH and GSH synthesis enzymes in response to the treatment (Supplementary Fig. 10A, B and Fig. 5C). Notably, the fraction of oxidized glutathione (GSSG) dramatically increased exclusively in KRAS-WT cells. This finding suggests that the increased GSH production proved insufficient to overcome the high oxidative stress induced by TrxR1 inhibition in KRAS-WT LC cells. (Supplementary Fig. 10B). To proof this concept, we showed that supplementation with cell permeable GSH (GSH-EE) rescues the effect of TRXR1 inhibition in KRAS-WT cells (Supplementary Figure 10C). Conversely, inhibition of GSH synthesis with buthioninesulfoximine (BSO) dramatically resensitizes KMLC cells to TRXR1 inhibition (Supplementary Fig. 10D)^8,14,56–58^. Even though auranofin and CS47 produced a similar effect within the same genotype, and the KRAS-WT and EGFR-MUT LC cell cohorts shared the greatest DEG overlap (Fig. 5D), what captured our attention is that *HMOX1* was the only DEG upregulated in all KRAS-WT, EGFR-MUT and KM lines regardless of CS47 and auranofin treatment (Fig. 5E), and was also the most upregulated protein in sensitive cells (Supplementary Fig. 9). *HMOX1* encodes for heme oxygenase 1 (HO-1), a stress inducible enzyme that catalyzes the catabolism of heme using NADPH–cytochrome P450 reductase and oxygen, to generate biliverdin, ferrous iron (Fe^2+^) and carbon monoxide (CO) (Fig. 5F). HO-1 is usually expressed at very low level in most tissues but gets promptly transcriptionally upregulated under oxidative stress^26,59^. If a moderate upregulation of *HMOX1* can function as antioxidant, its excessive levels can produce iron overload and ferroptosis^26^. *HMOX1* along with *GCLM, GCLC, TXNRD1, NQO1* are *bona fide* targets of the nuclear factor-erythroid factor 2-related factor 2 (NRF2) that is activated by oncogenes like KM to lower ROS and promote tumorigenesis^60^. However, along with its antioxidant, pro-tumorigenic role, NRF2 hyperactivation has been also reported to be deleterious in a subset of lung cancer cells by inducing reductive stress^61^. Therefore, we tested whether NRF2 hyperactivation might be responsible for the lethal phenotype observed in KRAS-WT lung cancer cells. However, NRF2 activation with tert-Butylhydroquinone (TBHQ) or the KEAP1-inhibitor KI696, antagonizes with TRXR1 inhibition rather than synergizing with it in KRAS-WT LC cells (Supplementary Fig. 11A). This finding is consistent with previous studies showing that KEAP1 mutant cells are resistant to auranofin^57^ and suggests that NRF2 activation represents a classical antioxidant mechanism in response to TRXR1 inhibition. This observation is supported by DepMap data showing that *TXNRD1* and *NRF2* are anti-correlated in LC (Supplementary Fig. 11B, C). TBHQ or KI696 single treatments had no significant effect on KRAS-WT LC cells. Importantly, even though they induced targets like GCLM, GCLC, TRXR1, and NQO1, they did not upregulate HO-1 (Supplementary Fig. 11D). This evidence indicates that NRF2 activation alone is not sufficient to drive ferroptosis in KRAS-WT cells and support the hypothesis that HO-1 acts as a critical downstream effector of TRXR1 inhibition in this context.

**Figure 5.**
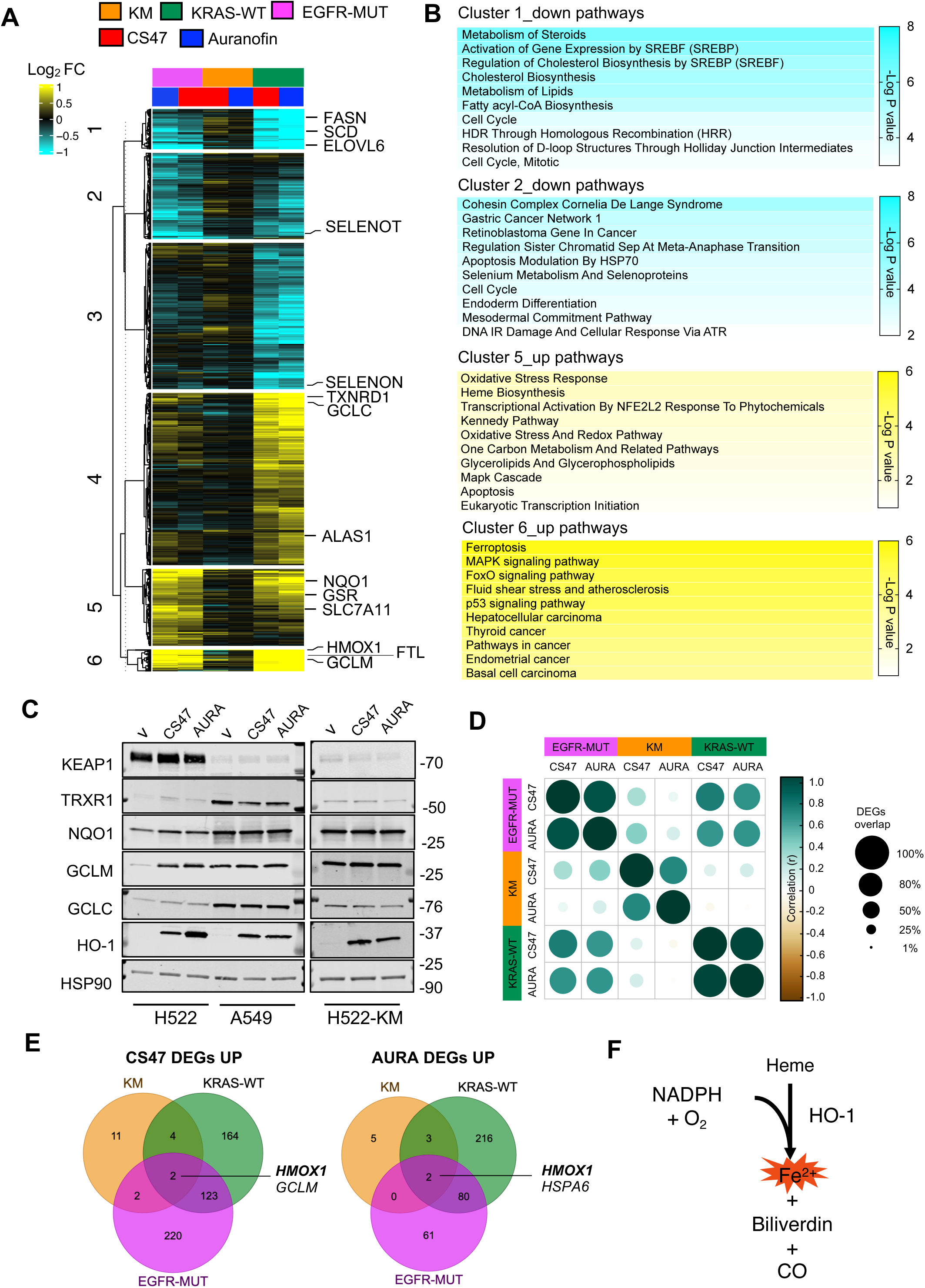
TRXR1 inhibition transcriptionally reprograms lipid, GSH and iron metabolism. A) RNA-seq heatmap showing differentially expressed genes (DEGs) in KM (H157, H1792, H460, A549), KRAS-WT (H522, H661, H1299, H1993) and EGFR-MUT (PC9, H1650, HCC827) LC cell lines treated as indicated for 24 hours. Values are reported as the average Log_2_ FC compared to the vehicle (0.5% DMSO) per each treatment and genotype group. B) Enrichr analysis showing the top pathways for the indicated clusters reported in panel A. C) Immunoblot validation of selected RNA-seq targets in the indicated cell lines following the specified treatments for 24 hours (n=2 independent experiments). D) Bubble plot comparing expression of DEGs across the various treatments and genotypes. Bubble size is proportional to size of the overlap, while color denotes Pearson correlation (r). E) Venn diagrams showing the common significant upregulated DEGs in all genotypes upon CS47 and auranofin treatments. F) Schematic of HO-1 enzymatic function.

### HMOX1 is necessary to drive ferroptosis under TXNRD1 inhibition in KRAS-WT lung cancer

To test this hypothesis, we ectopically expressed *HMOX1-GFP* in H522 and H661 KRAS-WT LC cells and sorted them for low (lo) and high (hi) GFP (Fig. 6A and Supplementary Fig. 12A). Upon treatment with CS47 or auranofin, HO-1 overexpression synergized with TRXR1 inhibition in a dose-dependent manner in both 2D and 3D culture systems (Fig. 6B-D and Supplementary Fig. 12B). A similar effect was achieved by pharmacological induction of HO-1 with Cobaltic Protoporphyrin IX Chloride (CoPP)^62,63^ in combination with CS47 (Supplementary Fig. 12C). We observed that *HMOX1-GFP^lo^*was sufficient to elevate intracellular Fe^2+^ levels beyond those observed with TRXR1 inhibition, consistent with TRXR1 inhibition inducing *HMOX1* expression and HO-1 enzymatic activity (Fig. 5 and Supplementary Fig. 12D, E). Conversely, inducible knock-down of *HMOX1* completely abrogated CS47-induced ferroptosis in the sensitive KRAS-WT and EGFR-MUT LC cells (Supplementary Fig. 12F-H), demonstrating that HO-1 is necessary to execute ferroptosis in this context. To validate these results *in vivo*, we established xenografts using parental or *HMOX1-GFP^lo^* H661 cells and treated mice with CS47 or vehicle (Fig. 6E). Notably, HO-1 overexpression alone delayed tumor growth so substantially that we postponed CS47 treatment by one week to allow tumor sizes to reach comparable baseline levels. While CS47 significantly inhibited tumor growth in parental H661 xenografts, its effect was even more pronounced in *HMOX1-GFP^lo^* tumors (Fig. 6E). This enhanced efficacy correlated with increased lipid peroxidation, as evidenced by the intratumor accumulation of the ferroptosis marker 4-Hydroxynonenal^64,65^ (4-HNE) (Fig. 6F).

**Figure 6.**
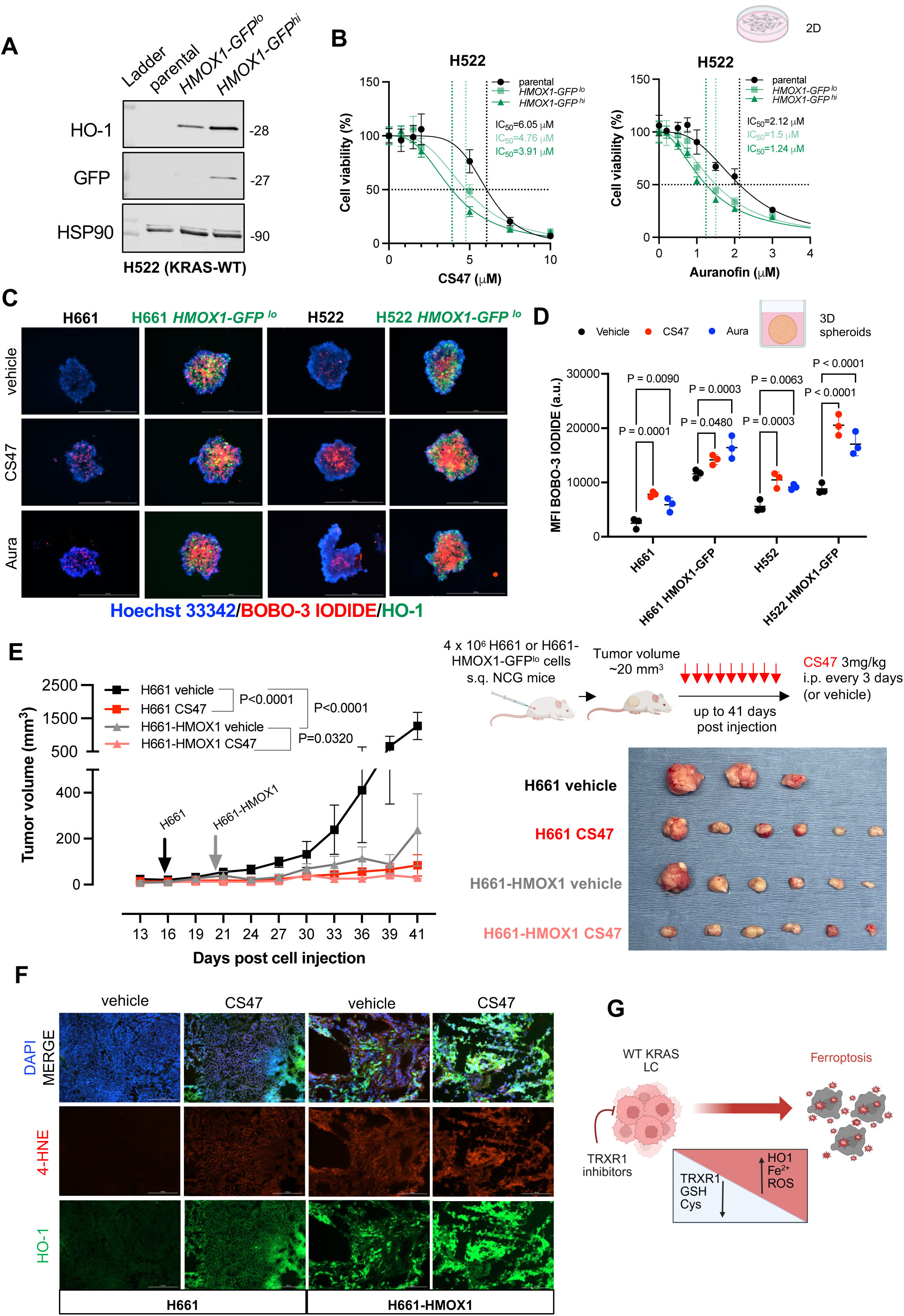
HMOX1 enables ferroptosis triggered by TRXR1 inhibition in KRAS-WT lung cancer. A) Immunoblot of H522 cells stably transfected and sorted for high (hi) and low expression (lo) of HMOX1-GFP (n=2 independent experiments). B) MTT viability assay and IC50 values of the indicated cell lines treated with CS47 and auranofin for 48 hours. C) Representative images and (D) quantification of Live/Dead assay in the indicated spheroids treated with vehicle, CS47 or auranofin for 48 hours (n= 3 independent replicates). P values were calculated using one-way ANOVA followed by Sidak’s tests. E) Tumor growth curves, schematic and post-dissection images of H661 and H661-HMOX1-GFP^lo^ xenografts (n=3-6 mice/group) treated as indicated. Arrows indicate treatment start. P values represent two-way ANOVA followed by Tukey’s multiple comparisons at 41 days. F) 4-HNE and HO-1staining in the indicated xenograft tumors (n=3 mice/group). G) Model of action by which TRXR1 inhibition depletes antioxidants GSH, Cysteine and induce HO-1, iron (Fe^2+^) and ROS overload in KRAS-WT LC.

Collectively, these results support a model in which TRXR1 inhibition, under conditions of low basal GSH leads to rapid ROS accumulation that drives HO-1 overexpression and intracellular iron overload, ultimately triggering ferroptosis in KRAS-WT LC cells (Fig. 6G).

### KMLC cells that become refractory to KRAS inhibition gain sensitivity to TRXR1 inhibitors

Our data indicate that an inverse relationship exists between KRAS dependency and TRXR1 sensitivity in LC, with KM signaling promoting resistance to TRXR1 inhibition (Fig. 1C, D; Fig. 2G and Supplementary Fig. 1C).

Mechanistically, this resistance is, in great part, driven by alternative antioxidant and lipid-remodeling pathways hijacked by KM cells to evade ferroptosis, including elevated GSH levels, enhanced fatty acid synthesis, and activation of the Lands Cycle^11^. KM is necessary and sufficient to induce this phenotype in LC, as we demonstrated in this paper and in our previous study^11^. Several KRAS inhibitors (KRASi), including sotorasib^66^, MRTX-1133^67^, RMC-7977^68^, RMC-6236^69,70^ are now in the clinic or clinical development. However, resistance to KRASi monotherapy remains a major challenge, emerging universally across all agents^71–80^. While some resistance mechanisms have been described, such as KRAS mutations or adaptive feedback loops, over 50% of cases remain unexplained, suggesting that additional, non-genetic, “off-target” mechanisms might be exploited for therapeutic gain.

In view of the mutually exclusive relationship outlined so far between KM and TRXR1, we hypothesized that LC cells which persist KRASi would rely on TRXR1 for their redox homeostasis. To this aim, we tested a panel of KMLC lines, harboring various *KRAS* mutations and co-mutations, with combinations of several KRAS and TRXR1 inhibitors for 4 days (Fig. 7A). Strikingly, across all combinations and cell lines, TRXR1 inhibition synergized with KRASi, suggesting that acute KM ablation reprograms KMLC cells toward TRXR1 dependency to mitigate ROS-induced cell death (Fig. 7A).

**Figure 7.**
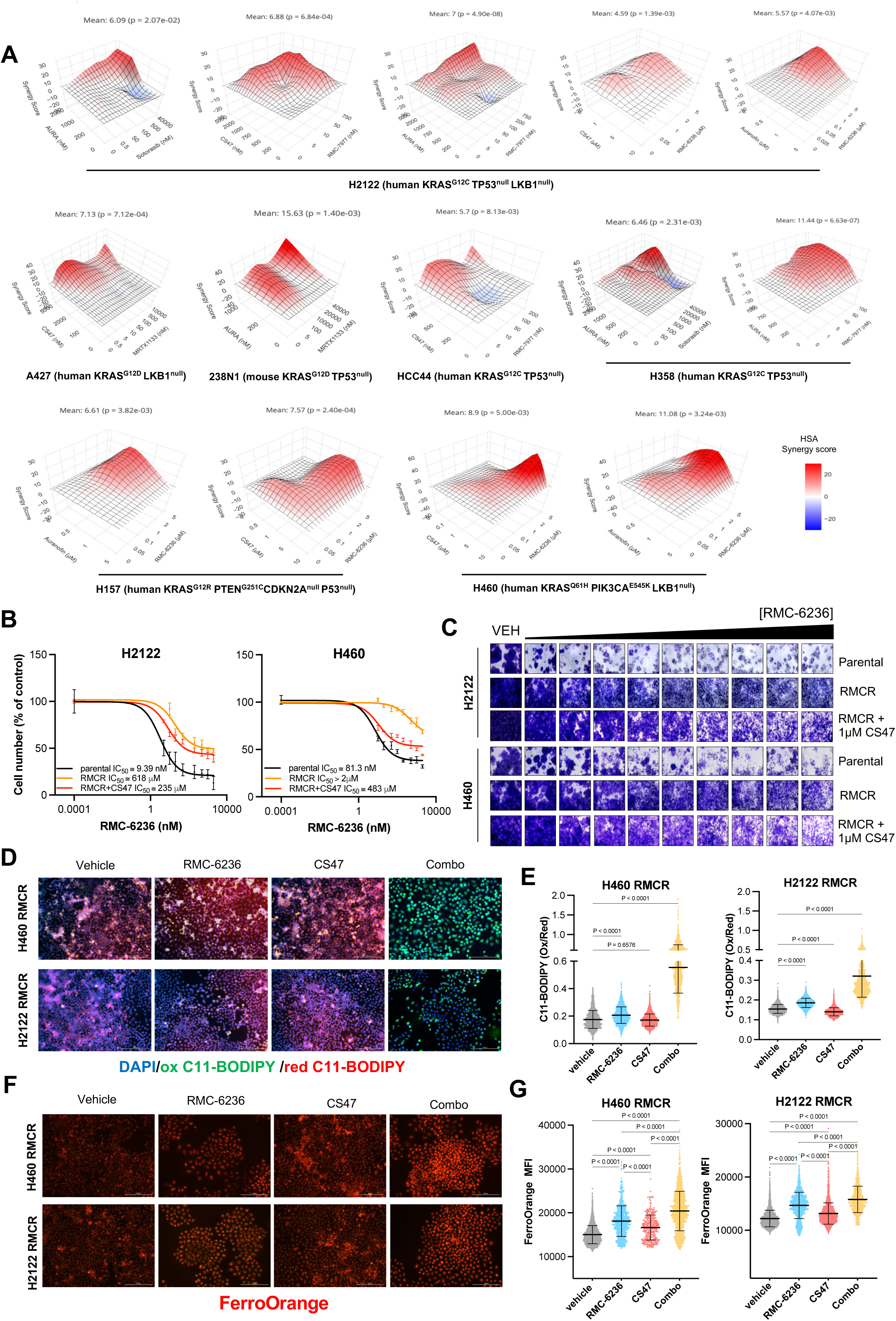
TRXR1 inhibition resensitizes LC persister cells to KRASi. A) Synergy plots of the indicated treatment combinations in the reported KM LC cell lines assessed as crystal violet assay at 4 days (n=2 independent experiments/each combination). P values for the average synergy scores were derived by bootstrapping of the dose–response matrix in www.synergyfinderplus.org. B) Curves and representative images (C) of crystal violet assays performed on H2122 and H460 parental and persister (RMCR) cells treated as indicated. Combination was done in presence of 1 µM CS47. Cells were maintained in 50 nM RMC-6236 for 3 weeks and then for 4 days as indicated. D) Representative images and (E) quantification of C11-BODIPY staining in H2122 and H460 RMCR cells treated as indicated (50 nM RMC-6236, 1 µM CS47, alone or in combo). Plots represent pooled cells from n= 3 replicates per group. P values were calculated using one-way ANOVA followed by Sidak’s tests. F) Representative images and (G) quantification of FerroOrange staining in H2122 and H460 RMCR cells treated as in D, E. Plots represent pooled cells from n= 3 replicates per group. P values were calculated using one-way ANOVA followed by Sidak’s tests.

To test whether this synergy results in ferroptosis, we used RMC-6236 (daraxonrasib), a first-in-class, oral RAS(ON) inhibitor targeting GTP-bound RAS^69,70^ which is now in Phase III clinical trials (NCT05379985, NCT06625320). After 3 weeks of RMC-6236 treatment, both H460 (*KRAS^Q61R^*, predicted to be less sensitive to RAS(ON) inhibition) and H2122 (*KRAS^G12C^,* among the most sensitive to RAS(ON) inhibition) cells developed a drug-persistent phenotype with significantly increased IC_50_ values (Fig. 7B, C). However, when co-treated with CS47, these KRASi-resistant (RMCR) cells underwent pronounced ferroptosis, evidenced by elevated C11-BODIPY oxidation (Fig. 7D, E) and intracellular Fe^2+^ accumulation (Fig. 7F, G). Notably, while both CS47 and RMC-6236 alone increased Fe^2+^ levels, only the combination robustly induced lipid peroxidation. This synergy was not due to enhanced ERK pathway inhibition. In fact, KRASi and TRXR1 co-treatment led to either increased or unchanged ERK phosphorylation despite inducing cell death (Supplementary Fig. 13). This finding is consistent with ROS-mediated ERK activation, previously described in other models of oxidative stress-induced cell death^21^. Rather, this ferroptosis switch was further supported by synergistic induction of HO-1 upon KRASi and TRXR1 co-treatment in both parental KM cells (4 days of treatment) and RMCR cells (Supplementary Fig. 13).

Altogether, these findings unveil that prolonged KRASi rewires LC cells toward a ferroptosis-prone state by promoting ROS and iron buildup. This redox imbalance creates a lethal window that can be fully exploited by TRXR1 inhibition, driving ferroptosis.

## Discussion

In cancer research, ferroptosis has emerged as an innate tumor suppressor mechanism that can be leveraged for therapeutic gain^8,11,81,82^. Yet, paradoxically, cancer cells often evolve to evade this fate, activating diverse anti-ferroptosis defenses to resist treatment and fuel metastatic progression ^9,11,83–86^. As these strategies can be tissue-, drug- and oncogene-dependent, it is critical to investigate the determinants of ferroptosis vulnerabilities in any given context^87^.

Our group previously demonstrated that KMLC cells evade ferroptosis by upregulating FASN activity and engaging the Lands cycle for lipid remodeling^11^. Crucially, KM is both necessary and sufficient to establish this ferroptosis-resistant phenotype in LC, as KRAS-WT, EGFR-mutant cells, or KRAS-mutant cells that had acquired resistance to KRASi all remained refractory to FASN inhibition-induced ferroptosis^11^. Even though these results were consistent with early evidence linking KRAS signaling and RAF/MEK/MAPK pathway to ferroptosis sensitivity^3,88–90^, they also raised a question: how do LC cells lacking an active KM survive in the same high-PUFA-PL- and oxygen-rich tumor niche without succumbing to ferroptosis?

Here, integrating DepMap data with functional assays, we revealed that LC cells that are KRAS-WT and/or KRAS independent are selectively vulnerable to TRXR1 inhibition. Notably, treatment with reversible TRXR1 inhibitors was sufficient to trigger ferroptosis, both *in vitro* and *in vivo*, showing less toxicity than for the FDA-approved auranofin. This property likely stems from our gold(I) compounds binding to an allosteric site akin to the recently described “Doorstop” pocket in *Schistosoma mansoni* thioredoxin glutathione reductase (TGR), a homolog of TRXR1^40^. This pocket, conserved in the human TRXR1, is distinct from the enzyme’s catalytic center and is defined by a unique amino acid composition that differentiates TRXR1 from closely related enzymes such as glutathione reductase (GSR). Notably, this binding mode allows our inhibitors to be specific for TRXR1 and to retain efficacy even in conditions of selenium-deprived TRXR1, where canonical TRXR1 inhibitors often fail^91^.

Mechanistically, ferroptosis requires the convergence of three key hallmarks: PUFA-PL, Fe^2+^ and ROS^92^. TRXR1 inhibition drives the accumulation of all three components, and ferroptosis can be effectively rescued by antioxidants (e.g., cysteine, NAC, GSH), lipid ROS scavengers (e.g., Lip-1, Fer-1), or iron chelation (e.g., DFO, or HMOX1 knockout) in sensitive KRAS-WT and EGFR-MUT LC cells. Conversely, as shown here and in previous work from our group and others, KM promotes resistance to TRXR1 in LC cells via multiple anti-ferroptosis mechanisms such as enhancing GSH biosynthesis^93^, remodeling phospholipids toward more oxidation-resistant species^11^, and potentially adapting to or co-opting Fe^2+^ accumulation, a known metabolic addiction in KM-driven cancers^14,94^. This would explain why previous studies done in KM tumor models concluded that TRXR1 is dispensable for tumorigenesis and cancer progression in the absence of other redox or metabolic perturbations^14,18^. Notably, under KRASi, we demonstrated that KMLC cells become susceptible to TRXR1 inhibition. Now that many KRASi are available and used in the clinic, our data offer an opportunity to further investigate whether TRXR1 inhibition can be leveraged to prevent or revert resistance to KRASi in patients and to deepen the knowledge about KM- and non-KM-specific ferroptosis regulation.

More broadly, in the context of therapy resistance across non-KM cancer types other than LC, progressive upregulation of *TXNRD1* and *HMOX1,* along with increasing dependence on *TXNRD1*, has been found to characterize persister cancer cells arising under various targeted therapies^95^. These findings further support the relevance of our observations in KRAS-WT and KRASi persister lung cancer cells and suggest that TRXR1 inhibition may offer a promising strategy to enhance therapeutic response in combination treatment approaches across multiple cancer contexts.

## Methods

### DepMap data

DepMap data including tumor types, mutational status, Genome-wide CRISPR knockout screens, and co-dependency correlations were accessed at https://depmap.org/portal/, release 23Q4 (https://doi.org/10.25452/figshare.plus.24667905.v2)^27,96–99^.

### Synthesis and stability of the gold(I) compounds

CS47 (4,5-dichloro-1H-imidazolate-1-yl)-(triphenylphosphane)-gold(I) and DM20 (4,5-dicyano-1H-imidazolate-1-yl)-(triphenylphosphane)-gold(I) were synthesized as previously described^31,32^. Briefly, the synthesis involves the reaction of the triphenylphosphine gold(I) tetrafluoroborate (PPh_3_Au⁺BF_4_⁻) salt with a tetrahydrofuran (THF) solution containing the appropriate sodium or potassium imidazolate. The resulting white solid is collected by evaporation to dryness, redissolved, filtered, and subsequently crystallized by slow diffusion of diethyl ether or hexane into methanol or chloroform solutions of the synthesized compounds.

The experimental protocol for the drug stability was adapted from Babak *et al*.^100^. In this approach, both ^1^H NMR spectroscopy and electrospray ionization mass spectrometry (ESI-MS) were employed. All chemicals were purchased from Merck and used without further purification. Solvents were obtained from Carlo Erba (Milan, Italy). Celite and molecular sieves were supplied by Supelco (Merck).

For the ^1^H NMR analysis, 15 mg of the gold compound was dissolved in 0.7 mL of DMSO-d_6_, and the solution was monitored over the course of one week to evaluate its stability and possible decomposition in a polar medium. ^1^H NMR spectra were recorded on an Oxford-400 Varian spectrometer operating at 400.4 MHz. Chemical shifts are reported in parts per million (ppm) relative to tetramethylsilane (Me_4_Si) as an internal standard.

For ESI-MS, the compounds were incubated in methanol diluted with an aqueous 10 mM ammonium acetate buffer (pH 7.4) at a 1:10 (v/v) ratio, resulting in a final concentration of 500 μM. The mixtures were incubated at 37°C, and aliquots were collected at 2 and 24 hours for analysis. ESI-MS spectra were acquired in both positive- and negative-ion modes using a Waters Alliance 2695 high-performance liquid chromatography (HPLC) system coupled to a Waters Micromass ZQ single quadrupole mass spectrometer. The mobile phase consisted of acetonitrile or methanol, and samples were prepared at a concentration of approximately 0.1 mM. A 1 µL injection volume was used, with a flow rate of 200 µL/min. Nitrogen served as both the drying and nebulizing gas. Capillary voltages were set to 4000 V for positive-ion mode and 3500 V for negative-ion mode.

### Cell culture

LC cell lines were originally obtained from the cell line repository of the Hamon Center for Therapeutic Oncology Research (UT Southwestern Medical Center)^101^. IMR-90 human lung fibroblasts were from ATCC (CC-186^TM^). The mouse 238N1 cells, previously generated from lymph nodes metastases of *Kras^LSL-G12D^*; *Trp53^f/f^* lung-adenocarcinoma-bearing mice^102^ were provided by Dr. Monte Winslow. H522-KM (*KRAS^G12D^)* were previously established in our lab^11^. All LC cells were maintained in HyClone RPMI medium supplemented with 10% heat-inactivated FBS and 1% penicillin-streptomycin, in humidified atmosphere, 5% CO_2_, 37°C. All cell lines were authenticated by DNA fingerprints for cell-line individualization using Promega Stem Elite ID system, a short tandem repeat (STR)-based assay and routinely confirmed to be *Mycoplasma* negative. For pharmacological treatments, LC and IMR-90 cells’ growth medium was replaced by RPMI medium supplemented with 5% heat-inactivated dialyzed FBS and 1% penicillin-streptomycin as previously described^11^.

### Plasmids, siRNAs and virus production

The shRNA inserts were cloned into the vector LT3GEPIR (Addgene, #111177) between the XhoI and EcoRI sites. The shRNAs target sequences (5’-3’) were from the library described by Feldmann et al.^103^: *shTXNRD1.2439* TATGATATTAATAACTTCCTTT; *shTXNRD1.328* TTATTTGTTGCCTTAATCCTGT; *shHMOX1.631* TAGAGCTGCTTGAACTTGGTGG; *shHMOX1.1537* TTTCACACAAAAGTTAGACCAA. Lentivirus production was carried out in HEK293T as previously described^104^.

pCX-HO1-2A-EGFP was a gift from Roberto Giovannoni (Addgene, #74672) and it was inserted into LC via electroporation using the NEPA21 electroporator (Bulldog-Bio) following the manufacturer’s guidelines. Stable transfected cells were obtained via G418 selection (1.5 mg/ml).

Predesigned *TXNRD1* siRNAs (#1, SASI_Hs02_00318195; #2, SASI_Hs02_00318197) and universal MISSION® siRNA Universal Negative Control #1 (cat. #SIC001) and #2 (cat. #SIC002) were from Millipore Sigma. DharmaFECT 4 (Thermo Scientific) was used for siRNA transfection.

### In silico docking

Human cytosolic TRXR1 (uniprot Q16881 pdb code 2CFY and 1W1C, isoform 5-major) was retrieved from Brookhaven Protein Data Bank (http://www.wwpdb.org). The 3D protein structure was optimized by AMBER force field (AMBER99SB-ILDN) within GROMACS 2024^105^ framework, and then they were used in the subsequent molecular docking calculations. The 3D structure of the gold compounds was retrieved from Cambridge Crystallographic database CSDD [ID 997733, 997734]. Ligands’ Mulliken charges were calculated at B3LYP/6-31G** using G016 software^106^. Autodock 4.2/ MGLTools1.5.7 was used to perform the molecular docking calculations^107^. For both compounds, a blind docking approach was used at first to identify by cluster analysis every putative site on the TRXR1 molecular surface. Subsequently, on the lowest energy and most populated poses, a focused docking protocol has been applied to better refine both pose and its energy. For the Blind docking, the grid map, centered in the center of mass of the enzymes was set to include the whole protein; in the focused docking protocol, the grid map was centered on the ligand and extended around the cleft (40 × 40 × 40 Å^3^) with points spaced equally at 0.375 Å. The number of GA (genetic algorithm) runs was set to 100, the energy evaluations (25 000 000), the maximum number of top individuals that automatically survive (0.1) and the step size for translation (0.2 Å). All the docking calculations were carried out in triplicate using three different CPUs random seed. The final docked ligand-protein complexes were ranked according to the predicted binding energy and all the conformations were processed using the built-in clustering analysis with a 2.0 Å cutoff. The lowest energy clusters were then used for subsequent analysis. Molecular graphics images were produced using the UCSF Chimera 1.17 package (Resource for Biocomputing, Visualization, and Informatics at the University of California, San Francisco, CA, USA). The ligand-enzyme interactions were analyzed by CHIMERA and PLIP (protein ligand interaction)^108,109^.

### Drugs and enzymatic inhibitors

auranofin (#15316), Ferrostatin-1 (Fer-1, #17729), Liproxstatin-1 (Lip-1, #17730), Deferoxamine (mesylate) (DFO, #14505), L-selenocystine (#17793), Cobaltic (III) protoporohirin IX chloride (CoPP, #33794), Glutathione ethyl ester (GSH-EE, #14953), and L-Buthionine-(S,R)-Sulfoximine (BSO, #14484) were from Cayman Chemicals. Sotorasib (#HY-114277), MRTX-1133 (#HY-134813), RMC-7977 (#HY-156498) and KI696 (#HY-101140) were from MedChemExpress. RMC-6236 (#207192) was from MedKoo Biosciences. tert-Butylhydroquinone (TBHQ, #112941), N-Acetyl-L-Cysteine (NAC, #A7250) and L-cysteine (#C7352) were from Millipore Sigma.

### 2D and 3D cell viability

For 2D cultures, 0.8-1 x 10^4^ cells per well were seeded into flat bottom 96-well plates one day prior to treatment to ensure approximately 60% confluence at the time of drug administration. Cells were treated for either 2 days (for compounds such as CS47, DM20, and auranofin alone or in rescue experiments) or 4 days (for all KRASi alone or in combination). At the treatment endpoint, cell viability was assessed via MTT or Crystal Violet Assay as previously described^11^.

For MTT assay, 10 μL of MTT solution (5 mg/mL) was added to each well. After a 4-hour incubation at 37 °C, the medium was removed, and formazan crystals were dissolved in 200 μL of DMSO per well. Plates were incubated for 10 minutes at 37 °C, and absorbance was measured at 570 nm (OD 570 nm). For Crystal Violet assay, the medium was removed and wells washed twice with PBS before incubation with 0.5% Crystal Violet Staining Solution (in 20% v/v methanol) for 20 minutes at RT. Plates were then washed at least three times in tap water and let air dry. Pictures were taken and confluence quantified using Lionheart FX (Agilent) and Gen5 software (v3.15). For combination treatments, synergy plots were made using SynergyFinder Plus (www.synergyfinderplus.org)^110^.

For 3D cultures Live/Dead assay, 2-3 x 10^3^ cells/50 µl per well were seeded into U bottom ultra-low attachments Nunclon Sphera 96-well plates (Thermo Scientific, #174925) two days prior to treatment to allow spheroid formation. Treatments were added (50 µl per well) as described for 2D conditions. At endpoint, 40 μL of media was carefully removed from the wells and replaced with serum- and phenol-red free OptiMEM containing NucBlue Live Cell Reagent (Hoechst 33342, Invitrogen, #R37605) and 0.5 µM BOBO-3 Iodide (Invitrogen, #B3586). Plates were incubated for 30 minutes in humidified atmosphere, 5% CO_2_, 37°C. Images were acquired and quantified using Lionheart FX (Agilent) and Gen5 software (v3.15).

### Mouse studies

All animal studies were approved by the Institutional Animal Care and Use Committee (IACUC) at the University of Cincinnati (protocol 24-01-05-01) and UT Southwestern Medical Center. For tumor studies power calculation was conducted using ClinCalc.com using mean tumor burden from previous mouse experiments^11^ and both male and female were included in the analysis.

#### LC-MS/MS Conditions for Pharmacokinetic (PK) Study

CS47 was chelated by exchange with diethyldithiocarbamate (DDTC) for detection by LC-MS/MS for *in vivo* analysis. Briefly, Gender Pooled CD1 mouse plasma (MSE00PLK2-470804) from BioIVT (Westbury, NY) was spiked with varying amounts of CS47 dissolved in DMSO (standards) or were obtained from a PK study (described below). An aliquot of 10 uL of each sample was diluted with 90 uL of PBS. Then 30 uL of a solution containing DDTC (20 mg/mL in 0.1 M NaOH), 50 ng tolbutamide, and 200 ng Bis(triphenylphosphine)palladium(II) dichloride ((NH_3_)_2_PdCl_2_) was added to each sample. The mixtures were vortexed vigorously for 1 min followed by incubation at 42°C for 40 min. Samples were extracted with 1 ml of acetonitrile and after vortexing and centrifugation, the supernatant was collected and dried under vacuum. Samples were reconstituted in 80% acetonitrile/20% water, centrifuged to remove any particulate material and evaluated by LC-MS/MS using an AB/Sciex (Applied Biosystems, Foster City, CA) TripleQuad™ 4500 mass spectrometer coupled to a Shimadzu (Columbia, MD) Prominence LC. The compound was detected using electrospray ionization with the mass spectrometer in positive MRM (multiple reaction monitoring) mode by following the precursor to fragment ion transition 492.9 (M + H^+^) → 116. Instrument settings were as follows: Dwell time of 100 ms, DP (declustering potential) 1.0 volts, EP (entrance potential) 10 volts, CE (collision energy) 39 volts, CXP (collision cell exit potential) 18 volts, CUR (curtain gas) 25, CAD (collision gas) 10 psi, IS (ion spray voltage) 4000 V, TEM (turbo heater temperature) 650°C, GS1 (nebulizing gas) 60 psi, GS2 (auxiliary gas) 60 psi. An Agilent (Santa Clara, CA) Poroshell 120 EC-C18 column (50 X 3 mm, 2.7 micron packing) was used for chromatography with the following conditions: Buffer A: dH20, Buffer B: acetonitrile; 0-1 min 5% B, 1-2.5 min gradient to 100% B, 2.5-7 min 100% B, 7-7.1 min gradient to 5% B, 7.1-7.5 min 5% B. Flow rate: 0.8 mL/min. Tolbutamide (transition 271.2 (M + H^+^) → 91.2) was used as an internal standard for LC-MS/MS quantitation, and chelation of ((NH_3_)_2_PdCl_2_) was monitored as a qualitative control for efficiency of exchange with DDTC. Uniform conversion was observed in all samples and standards.

#### Toxicity Testing

CS47 was prepared for dosing by dissolving the compound in DMSO and diluting to a final mixture of 5% DMSO/5% Cremophor EL/90% D5W. Female Nod Scid IL-2Rψ^-/-^ (NCG, strain #572) mice at 9 weeks of age (Charles River, Wilmington, MA) were dosed in duplicate a single time i.p. with various concentrations of CS47 (3, 5 and 10 mg/kg) in comparison to vehicle only. Blood was collected from each animal at 24 hours. Animals were observed for one month following dosing.

#### Pharmacokinetic Study

NCG mice at 9 weeks of age (strain #572, Charles River) were dosed IP with CS47 formulated as above at 10 mg/kg (10 ml/kg) as acutely the compound did not show significant effects at this dose. Blood was collected into K_2_EDTA tubes after various times after compound administration. The blood was subsequently centrifuged at 9300 x g for 10’ to isolate plasma. Plasma was stored at -80°C until analysis. Samples were processed as described above. Quantitation was performed directly in Analyst 1.7.3. A value of 3X above the signal obtained from blank plasma or tissue was designated the limit of detection (LOD). The limit of quantitation (LOQ) was defined as the lowest concentration at which back calculation yielded a concentration within 20% of theoretical. The LOQ for CS47 was 5 ng/ml in plasma. In general back calculation of points on both curves yielded values within 15% of theoretical over four orders of magnitude (5000 to 5 ng/ml). A standard noncompartmental analysis with sparse sampling was performed for initial data evaluation using CS47 concentration-time profiles. The maximum plasma drug concentration (C_max_) and time to reach C_max_ (T_max_) were obtained directly from the experimental data. Terminal half-life, area under the concentration time curve from time zero to the last measured timepoint (AUC, calculated by linear trapezoidal method) were calculated using the noncompartmental analysis tool in Phoenix™ WinNonlin® (Pharsight, Mountain View, CA).

### 4-HNE staining

Tumor cryosections (10 μm thick) were prepared using a cryostat and air-dried for 30 minutes at room temperature. Sections were fixed in 4% paraformaldehyde (PFA) in PBS (pH 7.4) for 20 minutes, then washed with PBS and permeabilized with 0.2% Triton X-100 for 20 minutes at room temperature. Blocking was performed with 2.5% horse serum (#S-2012-50, Vector Laboratories) for 30 minutes.

Sections were incubated overnight at 4°C in a humidified chamber with primary antibodies diluted 1:50 in 1% horse serum in PBS: either anti-4-HNE (Bioss, #BS-6313R) or anti-HMOX1 (Santa Cruz, #136960). Following three PBS washes, sections were incubated with Alexa Fluor 568 donkey anti-rabbit IgG (Invitrogen, #A10042) or Alexa Fluor 488 goat anti-mouse IgG (Invitrogen, #A11001) for 1 hour at room temperature. After additional PBS washes, nuclei were counterstained with DAPI, and slides were mounted using Fluoromount-G (Thermo Scientific). Imaging was performed using the Lionheart FX microscope (BioTek-Agilent), and analysis was conducted using Gen5 software (v3.15).

### TRXR1 activity

3 x 10^5^ LC cells per well were plated onto 6-well plates and a day after were treated with CS47 (2 μM), DM20 (2 μM), auranofin (1 μM) for 24 hours. Cell pellets were lysed in passive lysis buffer Passive Lysis 5x Buffer (Promega, #E1941) diluted 1:5 supplemented with TRXR1 activity was measured in the cell lysates using both the Thioredoxin Reductase Assay Kit (Abcam, #ab83463) as previously described^29^ and a continuous direct enzymatic assay adapted from Cunniff et al.^111^. This assay relies on the NADPH-dependent reduction of L-selenocystine, with activity monitored via the decrease in absorbance at 340 nm. Compared to other methods, this approach provides high specificity for TRXR1. The assay was optimized for buffer composition, pH, and temperature. The final reaction buffer contains 10 μM ascorbic acid in 40 mM Tris-HCl (pH 7.5) with 0.1 mM EDTA. Selenocystine (4 mM stock) was prepared by dissolving 1.6 mg selenocystine in 200 μL of 1 N NaOH, followed by addition of 100 μL of 1 N HCl and 700 μL of ddH₂O. A master mix was prepared containing 1 mM NADPH, 2 mM selenocystine, 10 μM ascorbic acid, 24 mM Tris-HCl (pH 7.5), 0.2 mM EDTA. Reactions were carried out in 96-well clear round-bottom UV plates (Corning, #3635) using 40 μL of master mix and cell lysate diluted in ddH₂O to a final volume of 100 μL and final NADPH concentration of 400 μM. Absorbance at 340 nm was recorded kinetically for 20 minutes at 37 °C using a POLARstar Omega plate reader (BMG Labtech). Enzymatic activity was calculated using the extinction coefficient of NADPH (ε = 6220 M⁻¹ cm⁻¹) and normalized to total protein content as determined by Bradford assay (μmol NADPH/min/mg protein). Residual activity was expressed as a percentage relative to vehicle treated lysates.

### C11-BODIPY staining

Cells were seeded on coverslips and treated for 24 hours as indicated. Coverslips were incubated with 2.5 μM BODIPY 581/591 C11, washed with PBS, fixed with 10% formalin for 1 hour, counterstained with DAPI and mounted using Fluoromount-G medium (Thermo Scientific). For cells, three images per slide were acquired using a Zeiss LSM 710 confocal microscope equipped with a Plan-Apo 63x/1.4 oil DIC M27 objective and ZEN 2010 B SP1 software (Zeiss). For tumors, snap frozen tissues were equilibrated at -20°C for 30 min, prior to cryostat cut (10 μm slices). Tissue slices were equilibrated at RT (10 min) and incubated with 2.5 μM BODIPY 581/591 C11 in a humidified chamber. Slides were washed, fixed, and mounted as described above for cells. Three images per sample were acquired using Lionheart FX (Biotek-Agilent) and analyzed with Gen5 software (v3.15). For the ratiometric quantification of oxidized to reduced C11-BODIPY fluorescence (C11ox/C11red) per image or region of interest (ROI). Analysis was conducted using either Gen5 software or ImageJ (similar software can be used): 1) Apply a primary mask to DAPI-stained nuclei; 2) Expand a secondary mask identically on both Red and GFP channels; 3) Measure mean fluorescence intensity (MFI) for each channel and calculate C11ox/C11red ratio (GFP MFI/Red MFI).

### FerroOrange staining

Cells were seeded onto 24-well, flat glass bottom (Greiner, #662892) and treated as indicated for 24 hours. Cells were then washed 3 times with HBSS solution and then incubated with 1μM FerroOrange (Dojindo, #F374) at 37°C in the cell culture incubator, for 30 minutes. Images were then acquired using a Lionheart FX (BioTek) and analyzed with Gen5 software (v3.15). Texas Red (EX 586/15 nm, EM 647/57 nm) MFI was calculated per each cell in a total of 3 replicates per condition.

### MDA detection

MDA was quantified using the Lipid Peroxidation (MDA) Assay Kit (Sigma, #MAK085). In this kit MDA reacts with TBA to form a fluorogenic adduct detectable at an excitation wavelength of 532 nm and emission at 553 nm. Standard curves were generated using MDA standards at 0, 2, 4, 6, 8, and 10 μM in water. Cells treated with CS47 (2 μM), DM20 (2 μM), auranofin (1 μM) for 24 hours were lysed in 300 μL of lysis buffer with 3 μL of butylhydroxytoluene (BHT) to prevent further oxidation. Lysates were centrifuged at 13,000 × g for 10 min, and 200 μL of the supernatant were transferred to new tubes. Then, 600 μL of TBA reagent were added to each sample and standard. The mixtures were incubated at 95 °C for 60 min, cooled on ice for 10 min, and brought to room temperature. Fluorescence was measured using a PerkinElmer LS 55 spectrofluorometer. All samples were assayed in triplicate.

### Protein extraction and immunoblot

Following the indicated treatments, protein lysates were extracted using RIPA buffer (1% sodium deoxycholate, 150 mM NaCl, 1 mM EGTA, 1% NP-40, 20 mM Tris-HCl) supplemented with cOmplete, EDTA-free Protease Inhibitor Cocktail (Millipore-Sigma, #11836170001) and PhosSTOP (Millipore-Sigma, #4906845001). Protein concentration was assessed using Bio-Rad Protein Assay Dye Reagent Concentrate (Bio-Rad, #5000006). Samples (15-30 μg of total proteins) were prepared in 4X Laemli Sample Buffer (Bio-Rad, #1610747) supplemented with ß-mercaptoethanol, resolved on 4–15% Criterion TGX Precast Midi Protein Gels (Bio-Rad, #5671084) and transferred onto Amersham Protran 0.2 μm nitrocellulose membrane (Millipore-Sigma, #GE10600012). After blocking in 5% Blotting-Grade Blocker (Bio-Rad, #1706404) in 0.01% Tween, TBS buffer (TBS-T) for 1 hour at RT, membranes were incubated with primary antibodies overnight at 4°C. After washing (3 times) in TBS-T, membranes were incubated the appropriate secondary antibody for 1 hour at RT. The blots washed 2 times TBS-T, 1 time time in TBS and then acquired and analyzed using Odyssey Clx and Image Studio software v.6.0 (LICORbio). List of antibodies and dilutions as follows: anti-KEAP1 (Cell Signaling, #8047, 1:1000), anti-TRXR1 (Santa Cruz, #28321, 1:1000), anti-TRX1 (Santa Cruz, sc-271281, 1:500); anti-NQO1 (Boster Bio, # A00494, 1:1000); anti-GCLM (abcam, #ab126704, 1:1000); anti-GCLC (abcam, #ab190685, 1:1000); anti-HMOX1 (Santa Cruz, #136960, 1:500); anti-HSP90 (Cell Signaling, #4877, 1:1000); anti-GFP (Sigma, #G1544, 1:2000); anti-GAPDH (Santa Cruz, #32233, 1:500); anti-PRDX1 (abcam, #ab41906, 1:1000); anti-ERK (Cell Signaling, #4696, 1:2000), anti-pERK^T202/Y204^ (Cell Signaling, #9101, 1:1000); Donkey anti-mouse IgG (H+L) crossed-absorbed secondary antibody-DyLight 680 (Invitrogen #SA5-10170, dil 1:5000); Goat anti-rabbit IgG (H+L) crossed-absorbed secondary antibody-DyLight 800 (Invitrogen #SA5-10036, dil 1:5000); Goat anti-mouse IgG (H+L) crossed-absorbed secondary antibody-DyLight 800 (Invitrogen #SA5-10176, dil 1:5000).

The redox western blot for TRX1 and PRDX1 was carried out as previously described^112,113^. Briefly, after 6 hours of treatment with 1μM of auranofin or 2 μM of CS47, cell dishes were washed twice in PBS and incubated with 200 μL of freshly made alkylation buffer (40 mM HEPES, 50 mM NaCl, 1 mM EGTA, 25 mM N-ethylmaleimide) for 10 minutes at RT. Cell lysis was accomplished by adding 20 μL of 10% CHAPS to the dishes in alkylation buffer and incubating for additional 10 min at RT. Cell lysates centrifuged at 15,000 *g* for 20 min at 4°C, supernatants were isolated and quantified, and 15 μg of proteins resolved onto 4–15% Criterion TGX Precast Midi Protein Gels after mix with non-reducing 4X Laemli Sample Buffer.

### PISA assay and proteomics

#### Cell Culture and compound treatments

Cell line H522 were grown and expanded in RPMI medium + 10% fetal bovine serum + 1% penicillin/streptomycin. Cells were sub-cultured when 90% confluence was reached, and medium refreshed every 2/3 days. At 70% confluence, cells were washed twice in PBS 1X and treated with 1 µM and 5 µM CS47 or vehicle (DMSO) for PISA and 2 µM CS47 for Expression in RPMI medium supplemented with 5% dialyzed serum and 1% penicillin/streptomycin.

For each of the PISA assay with 1h treatment (4 replicates/condition) and Expression proteomics at 48h treatment (3 replicates/condition), cells were seeded at 10^6^/mL the day before the compound treatment into 16 x T25 flasks. Each compound was added from a stock solution of the compound in DMSO and equal volumes of DMSO were added for the control flasks (0.2%, final concentration). At the end of each treatment, cells were collected from each flask by centrifugation and washed twice in 10 ml of PBS. The cell pellet of each flask was resuspended into 1ml of PBS supplemented with protease inhibitors (Thermo Fisher) for PISA, or frozen as pellet at -80°C for Expression proteomics.

#### Sample preparation

In the PISA experiment, according to the published protocols^34,114,115^, cells were divided into 16 aliquots. Each aliquot was then incubated for 3 minutes at specific temperatures: 45.1, 46.5, 48.4, 50.3, 52.0, 53.9, 55.7, 57.1, 58.6, 60.0, 61.9, 63.8, 65.5, 67.4, 69.2, or 70.6 °C using a gradient thermal cycler (Eppendorf, Mastercycler X50s). After incubation, the samples were allowed to cool down at RT for 5 minutes and snap frozen into liquid nitrogen. Samples were then subjected to 5 cycles of freeze/thaw cycles between liquid nitrogen and 37°C incubation, all temperature pointś tubes for each replicate was pooled and incubated with 0.4% NP40 at 4°C for 1 hour, then ultra-centrifuged at 125,000 *g* at 4 °C for 30 min to separate the soluble fraction from the precipitated protein pellet.

In the expression proteomics experiment, cell pellets were resuspended in 250 µl of RIPA buffer supplemented with protease inhibitors, then lysed by 5 cycles of freeze/thaw cycles between liquid nitrogen and 37°C incubation, and probe sonicated for 1 min on ice (03 pulse, 03 pause, 10 times -30% amplitude). Protein concentration of the soluble fraction of each PISA sample and each cell lysate the expression proteomics experiment was measured by the micro-BCA kit (Thermo Fisher). 50µg of each sample was reduced with 8 mM DTT (Sigma), then alkylated with 25 mM IAA (Sigma). Samples were precipitated using 5 volumes of cold acetone at -20 °C overnight, digested by Lys C (Wako Chemicals GmbH) at a 1:75 enzyme to protein ratio for 6 hours at 30°C and then by trypsin (Promega) (1:50 enzyme to protein ratio) overnight. After labeling using 18 TMTpro reagents (Thermo Fisher Scientific) according to manufacturer’s instructions, each 18-plexed peptide samples of PISA and Expression proteomics were desalted with C18 Sep-pak columns (Waters). Peptides samples were then separated by high pH reversed phase chromatography using a Dionex Ultimate 3000 UPLC system (Thermo Fisher Scientific) at a 200 µl/min flowrate and finally collected into 48 fractions of 200. Each fraction was speed-vacuum dried, then resuspended in 0.1% FA, 2% ACN buffer and analyzed by nanoLC-MS/MS.

### NanoLC-MS/MS analysis

#### Data analysis

Proteomics data were analyzed using Proteome Discoverer against the Uniprot Homo sapiens database (UP000005640). Fixed modifications included cysteine carbamidomethylation and TMT tags; variable modifications included methionine deamination and oxidation on arginine/asparagine. Trypsin digestion allowed up to two missed cleavages. A 1% FDR was applied at protein and peptide levels. Contaminants and reversed sequences were removed, and only proteins with ≥2 unique peptides were quantified. For PISA (1 h) and expression (6 h) datasets, proteins with missing values across all replicates were excluded. Protein intensities were normalized to total protein per sample, and compound-to-control ratios were calculated and log2-transformed. Two-tailed t-tests were used to determine p-values. Volcano plots were generated, and targets were defined as proteins with high absolute log2 fold changes in the expression dataset and low p-values in the PISA dataset.

### Cellular Thermal Shift Assay (CETSA)

CETSA was performed in intact H522 cells following the protocol described by Molina et al. ^33^. On Day 1, ten 100-mm dishes were seeded with 2.4 × 10^6^ cells each to obtain sufficient material for a 10-temperature thermal gradient in quadruplicate per treatment condition.

On Day 2, cells were treated with 2 μM CS47 or vehicle (DMSO) for 3 hours. Following treatment, cells were harvested, counted (∼6 × 10^5^ cells per temperature replicate), and resuspended in PBS containing protease inhibitors (50 μL per replicate). The suspension was aliquoted into 0.2 mL PCR tubes and exposed to a thermal gradient ranging from 37 to 80 °C for 3 minutes per temperature, followed by rapid cooling to 25 °C. After brief centrifugation to collect condensation, samples were transferred to clean 1.5 mL tubes. Samples were subjected to three freeze–thaw cycles using liquid nitrogen and a 37 °C water bath (3 min each), then centrifuged at 12,000 × g for 20 min at 4 °C. Supernatants were collected, mixed with 4X Laemmli Sample Buffer (Bio-Rad, #1610747) supplemented with β-mercaptoethanol, and denatured. Proteins were resolved on 4–15% Criterion TGX Precast Midi Protein Gels (Bio-Rad, #5671084) and transferred for immunoblotting with anti-TRXR1 antibody (Santa Cruz, #28321, 1:1000). TRXR1 levels were normalized to a loading control (HSP90) and compared between DMSO- and CS47-treated conditions.

### RNA-seq and bioinformatic analysis

Total RNA was extracted using the TRIzol Reagent from Invitrogen following the manufacturer’s instructions. Total RNA was purified using the NucleoSpin RNA Clean-up Kit (Macherey-Nagel). NEBNext Ultra II Directional RNA Library Prep kit was used for polyA RNA-seq library preparation.

RNA-seq reads were aligned to the mouse reference genome (hg38) using STAR (v2.7.1a)^116^. For each sample, a BAM file was generated containing both mapped and unmapped reads across splice junctions. Secondary alignments and multi-mapped reads were removed using in-house scripts, and only uniquely mapped reads were retained for downstream analyses. GENCODE annotation for hg38 (version 32) was used for reference-guided alignment and downstream quantification. Gene-level expression was quantified using featureCounts (v2.0.1)^117^, based on protein-coding genes from the annotation files. Quality control metrics were assessed MultiQC (v1.0)^118^.

Counts were normalized using counts per million reads (CPM). Genes without reads in either sample were removed. Differential expression analysis was performed in R using a linear model followed by a post-hoc multiple comparison using the R package emmeans. P-values were adjusted for multiple comparisons using the Benjamini-Hochberg correction (FDR<0.05). Genes were considered differentially expressed by FDR < 0.05 and log2 Fold Change (FC) > 0.3 Pathway analysis was carried out using Enrichr https://maayanlab.cloud/Enrichr/^119–121^.

### LC-MS lipidomics

Cell pellets and tissues were stored at −80 °C and extracted via a modified Bligh/Dyer liquid–liquid extraction as we previously described^11^. Lipid extracts were resuspended in dichloromethane/methanol/IPA (2:1:1) with 8 mM NH_4_F and SPLASH LipidoMix internal standards. Lipids were separated on an Acclaim C30 column (3 μm, 2.1 × 150 mm, Thermo) at 35 °C using a 250 μL/min flow rate and 10 μL injection. Solvent A: 10 mM ammonium formate in 60:40 ACN:water with 0.1% FA; Solvent B: 10 mM ammonium formate in 90:10 IPA:ACN with 0.1% FA. A gradient from 50% A (3–50%) was applied.

Data were acquired on an Orbitrap mass spectrometer (Thermo Scientific) using HESI in both positive (3.5 kV) and negative (2.4 kV) modes. Capillary and heater temperatures were 350 °C and 275 °C, respectively; sheath and auxiliary gas at 45 and 8 units. Full scan (m/z 250–1200) was performed at 30,000 resolution with AGC target 2×10^5^ and 100 ms max injection time. NCEs of 25, 30, and 35% were used for fragmentation. Lipid identification and quantification were performed using LipidSearch 4.2 (Thermo Scientific). Tolerances: parent m/z 5 ppm, product m/z 10 ppm, quant m/z 5 ppm, RT 1 min. Allowed adducts: positive mode (+H, +NH_4_, +H-H_2_O, +2H); negative mode (−H, +HCOO, +CH_3_COO, −2H).

### MALDI-IMS lipidomics

Cryosections of 10 μm thickness were prepared using a cryostat and mounted onto indium tin oxide (ITO)-coated glass slides (Diamond Coatings, UK). The mounted sections were stored at −80 °C until use and dried for 30 min in a vacuum desiccator prior to matrix application. Optical images of the dried tissue sections were acquired using an Epson Perfection V850 Pro scanner (EPSON, Japan). A matrix solution of 2,5-dihydroxybenzoic acid (DHB, ≥98%, Sigma-Aldrich) was prepared at a concentration of 50 mg/mL in a methanol/water mixture (7:3, v/v) containing 0.1% trifluoroacetic acid (TFA), and applied using a TM-Sprayer (HTX Technologies LLC, USA) at a flow rate of 0.12 mL/min, a nozzle velocity of 1200 mm/min, a drying time of 2 s between passes, and a line spacing of 2.5 mm. Matrix was deposited in 22 passes at a nozzle temperature of 60 °C. Imaging mass spectrometry was performed using an UltrafleXtreme MALDI-TOF/TOF instrument (Bruker Daltonics, Germany) equipped with a SmartBeam II 1 kHz Nd:YAG laser (355 nm), operated in positive ion mode. External calibration was performed using red phosphorus standards covering an m/z range of 300-2000 Da. Data were acquired over an m/z range of 300–2000 with a laser focus set to ‘small,’ averaging 500 laser shots per pixel. Instrument parameters included a delay time of 180 ns, ion source 1 voltage of 25 kV, ion source 2 voltage of 21.65 kV, and a lens voltage of 9.2 kV. Spatial resolution was set to 100 μm. Raw data were processed using FlexImaging (v4.1), SCiLS Lab (v2023b), and METASPACE. Analysis included segmentation, identification of regions of interest (ROIs), spectral comparisons, and univariate statistical evaluation. Lipid annotations were assigned based on accurate mass measurements and validated against reference databases including LIPID MAPS (https://www.lipidmaps.org), SwissLipids (https://www.swisslipids.org), and METASPACE (https://metaspace2020.org). After MALDI-MS acquisition, the matrix was removed, and hematoxylin and eosin (H&E) staining was performed on the same tissue sections for histological evaluation.

### LC-MS metabolite detection

Sample preparation for the detection of steady-state cysteine, cystine, serine, glycine and GSH was performed following a previously established protocol^122^.

Immediately prior to metabolite extraction, a fresh NEM-containing extraction buffer was prepared by combining LC-MS grade methanol with 10 mM ammonium formate containing N-ethylmaleimide (NEM), along with internal standards: 10 μM ^13^C_3_,^15^N-NEM-cysteine, 4.18 μM 2H_5_-NEM-GSH, and 10 μL of Metabolomics Amino Acid Mix Standard (Cambridge Isotope Laboratories). Plates were transferred to ice, media was aspirated, and wells were washed with 1 mL of ice-cold PBS. Subsequently, 500 μL of chilled NEM-containing buffer was added to each well, incubated for 30 minutes at 4 °C, and cells were scraped and collected into 1.5 mL tubes. Samples were centrifuged at 17,000 × g for 20 minutes at 4 °C, and the resulting supernatants were stored at −80 °C until LC-MS analysis.

Metabolite extracts (50 μL) were transferred to LC-MS vials with inserts and analyzed using a Vanquish UPLC system coupled to a Q Exactive HF mass spectrometer equipped with a Heated Electrospray Ionization (HESI-II) source. Separation was achieved using an Atlantis Premier BEH Z-HILIC 2.5 μm 2.1 × 150 mm column maintained at 30°C. Mobile phase A consisted of 10 mM (NH_4_)_2_CO and 0.05% NH_4_OH in water; mobile phase B was 100% ACN. The injection volume was 5 μL. The gradient ran from 80% to 20% B over 13 minutes, followed by 2 minutes at 20% B, and re-equilibration to 80% B for 5 minutes. Autosampler washing was performed using 10% methanol at 1 μL/s for 10seconds. Mass spectra were acquired in positive ion mode over an m/z range of 65–950, at a resolution of 120,000. Prior to sample analysis, two blank runs followed by one conditioning run (first sample in the sequence) were conducted to stabilize the system. The LC-MS metabolite peaks were manually annotated and integrated using EI-Maven (v0.10.0). The peak areas of target metabolites were normalized by the peak area of their stable isotope labeled internal standard as previously described^123^.

### GSH/GSSG assay

GSH and GSSG were measured using the Glutathione GSH/GSSG Assay Kit (Millipore Sigma, # MAK440) following the manufacturer instructions starting from 2 x 10^6^ cells/sample/2 independent replicates treated with vehicle (DMSO) 2 μM CS47 or 1 μM auranofin for 24 hours.

### Statistical analysis

Statistical analyses were conducted using Microsoft Excel for Mac (v16.34) and GraphPad Prism (v10, GraphPad Software, San Diego, CA, USA). Data are presented as mean ± SD from at least two biologically independent experiments, with sample sizes (n) detailed in the figure legends. Comparisons between two groups were performed using two-tailed unpaired Student’s t-tests. For comparisons involving more than two groups, one- or two-way ANOVA was applied, followed by Dunnett’s, Tukey’s, or Sidak’s post hoc tests, or multiple two-tailed t-tests with FDR correction for multiple comparisons.

## Data availability

In silico docking pdb files are available in the Supplemental Information. RNA-seq data were deposited into GEO with accession number GSM9059311. Further information and requests for resources and reagents should be directed to and will be fulfilled according to institutional rules by the lead contact C. Andreani.

## Code availability

Custom R codes and data to support the analysis, visualizations, functional enrichments are available at https://github.com/BioinformaticsMUSC/AndreaniEtAl_2025. Some of the visualizations were carried out using BioRender https://www.biorender.com/

## Acknowledgements

C.A. was supported by University of Cincinnati, the Harris Scholar Award, the Grant #IRG-23-1141524- 01-IRG from the American Cancer Society, and the Marlene Harris Ride Cincinnati Cancer Pilot Program. Preliminary experiments were funded by NIH/NCI R01 CA259845-01A1 and The LCS Foundation (Ohio) (P.P.S.). We thank the National Center for Advancing Translational Sciences of the NIH UL1TR001425 (C.B.), Dr. Stephen Macha and the UC Department of Chemistry for the support in establishing the MALDI-IMS platform. We thank Dr. Ken D. Greis, Dr. Wendy Haffey, Dr. Birgit Ehmer for technical support and core resources at UC. The Chemical proteomics core facility of the Karolinska Institutet (Chemistry I Division, MBB Department), also Unit of SciLifeLab and node of the Swedish National Infrastructure for Biological Mass Spectrometry (BioMS), provided full support in the experimental design, performance, and data analysis of the proteomic studies. Bioinformatic analysis was supported by CNDD Genomics and Bioinformatics Core at MUSC and NIH P20GM148302 (S.B.). The authors thank the Erasmus+ KA171/2022 –Project n. 2022-1-IT02-KA171-HED-000073309 for supporting mobility and collaborative efforts between University of Camerino and University of Cincinnati. We additionally acknowledge the resources and expertise of the UT Southwestern institutionally supported Preclinical Pharmacology Core for PK studies.

## Author contribution

C.A. designed the study, performed the main experiments and wrote the first draft of the manuscript; C.B. performed the lipidomic and MALDI-IMS analyses, and contributed to all the in vivo studies; M.M. generated all the plasmids and lentiviral particles; N.S., L.L., R. Galassi designed and synthesized the gold(I) compounds, and performed stability assays; A.M. performed additional in vitro rescue experiments; R. Galeazzi performed the in silico docking; G.M.D performed the LC-MS metabolomics;

J.K. and N.W. performed the PK and acute toxicity studies; S.B. and C.A. performed bioinformatic RNA-seq analysis; M.G. performed the PISA and proteomics experiments; P.P. and S.S.O. assisted with cell culture and immunoblot experiments; A.T.M. performed blinded pathology assessment on tissues; S.P. performed the TRXR activity assay and the detection of MDA. C.A and P.P.S supervised the study and revised the manuscript with comments from all authors.

## References

1 Siegel, R. L., Kratzer, T. B., Giaquinto, A. N., Sung, H. & Jemal, A. Cancer statistics, 2025. CA: A Cancer Journal for Clinicians 75, 10–45 (2025). 10.3322/caac.21871

2 Meyer, M.-L. et al. New promises and challenges in the treatment of advanced non-small-cell lung cancer. The Lancet 404, 803–822 (2024). 10.1016/S0140-6736(24)01029-8

3 Dixon, S. J. et al. Ferroptosis: an iron-dependent form of nonapoptotic cell death. Cell 149, 1060–1072 (2012). 10.1016/j.cell.2012.03.042

4 Zhou, Q. et al. Ferroptosis in cancer: from molecular mechanisms to therapeutic strategies. Signal Transduction and Targeted Therapy 9, 55–55 (2024). 10.1038/s41392-024-01769-5

5 Dixon, S. J. & Olzmann, J. A. in *Nature Reviews Molecular Cell Biology* Vol. 25 (2024).

6 Conrad, M. Ferroptosis: when metabolism meets cell death. Physiological Reviews 105, 651–706 (2024). 10.1152/physrev.00031.2024

7 Stockwell, B. R. Ferroptosis turns 10: Emerging mechanisms, physiological functions, and therapeutic applications. Cell 185, 2401–2421 (2022). 10.1016/j.cell.2022.06.003

8 Bebber, C. M. et al. Ferroptosis response segregates small cell lung cancer (SCLC) neuroendocrine subtypes. Nature Communications 12, 2048–2048 (2021). 10.1038/s41467-021-22336-4

9 Ubellacker, J. M. et al. Lymph protects metastasizing melanoma cells from ferroptosis. Nature 585, 113–118 (2020). 10.1038/s41586-020-2623-z

10 Rodriguez, R., Schreiber, S. L. & Conrad, M. Persister cancer cells: Iron addiction and vulnerability to ferroptosis. Molecular Cell 82, 728–740 (2022). 10.1016/j.molcel.2021.12.001

11 Bartolacci, C. et al. Targeting de novo lipogenesis and the Lands cycle induces ferroptosis in KRAS-mutant lung cancer. Nature Communications 13, 4327–4327 (2022). 10.1038/s41467-022-31963-4

12 Zou, Y. et al. Plasticity of ether lipids promotes ferroptosis susceptibility and evasion. Nature 585, 603–608 (2020). 10.1038/s41586-020-2732-8

13 Mandal, P. K. et al. System x(c)- and thioredoxin reductase 1 cooperatively rescue glutathione deficiency. The Journal of biological chemistry 285, 22244–22253 (2010). 10.1074/jbc.M110.121327

14 Mandal, P. K. et al. Loss of Thioredoxin Reductase 1 Renders Tumors Highly Susceptible to Pharmacologic Glutathione Deprivation. Cancer Research 70, 9505–9514 (2010). 10.1158/0008-5472.CAN-10-1509

15 Conrad, M., Bornkamm, G. W. & Brielmeier, M. (eds Dolph L. Hatfield, Marla J. Berry, & Vadim N. Gladyshev) 195–206 (Springer US, 2006).

16 Asantewaa, G. et al. Glutathione synthesis in the mouse liver supports lipid abundance through NRF2 repression. Nature Communications 15, 6152 (2024). 10.1038/s41467-024-50454-2

17 Yan, X. et al. Inhibition of Thioredoxin/Thioredoxin Reductase Induces Synthetic Lethality in Lung Cancers with Compromised Glutathione Homeostasis. Cancer Research 79, 125–132 (2019). 10.1158/0008-5472.CAN-18-1938

18 Sherwood, A. M., Yasseen, B. A., DeBlasi, J. M., Caldwell, S. & DeNicola, G. M. Distinct roles for the thioredoxin and glutathione antioxidant systems in Nrf2-Mediated lung tumor initiation and progression. Redox Biology 83, 103653 (2025). 10.1016/j.redox.2025.103653

19 Shcholok, T. & Eftekharpour, E. Insights into the Multifaceted Roles of Thioredoxin-1 System: Exploring Knockout Murine Models. Biology 13 (2024).

20 Arnér, E. S. J. in Selenium and Selenoproteins in Cancer Vol. 136 (eds Kenneth D. Tew & Francesco B. T. Advances in Cancer Research Galli) 139–151 (Academic Press, 2017).

21 Akhiani, A. A. et al. Role of the ERK Pathway for Oxidant-Induced Parthanatos in Human Lymphocytes. PLOS ONE 9, e89646 (2014). 10.1371/journal.pone.0089646

22 Jakupoglu, C. et al. Cytoplasmic Thioredoxin Reductase Is Essential for Embryogenesis but Dispensable for Cardiac Development. Molecular and Cellular Biology 25, 1980–1988 (2005). 10.1128/MCB.25.5.1980-1988.2005

23 Gencheva, R. & Arnér, E. S. J. Thioredoxin Reductase Inhibition for Cancer Therapy. Annual Review of Pharmacology and Toxicology 62, 177–196 (2022). 10.1146/annurev-pharmtox-052220-102509

24 Soerensen, J. et al. The Role of Thioredoxin Reductases in Brain Development. PLOS ONE 3, e1813 (2008). 10.1371/journal.pone.0001813

25 Patwardhan, R. S., Rai, A., Sharma, D., Sandur, S. K. & Patwardhan, S. Txnrd1 as a prognosticator for recurrence, metastasis and response to neoadjuvant chemotherapy and radiotherapy in breast cancer patients. Heliyon 10, e27011–e27011 (2024). 10.1016/j.heliyon.2024.e27011

26 Hassannia, B., Vandenabeele, P. & Vanden Berghe, T. in *Cancer Cell* (2019). DepMap. (Broad, 2023).

27 Zheng, J. & Conrad, M. The Metabolic Underpinnings of Ferroptosis. Cell Metabolism 32, 920–937 (2020). 10.1016/j.cmet.2020.10.011

28 Galassi, R. et al. Anticancer Activity of Imidazolyl Gold (I/III) Compounds in Non-Small Cell Lung Cancer Cell Lines. Pharmaceuticals 17, 1133–1133 (2024).

29 Gambini, V. et al. In vitro and in vivo studies of gold(I) azolate/phosphane complexes for the treatment of basal like breast cancer. European Journal of Medicinal Chemistry 155, 418–427 (2018). 10.1016/j.ejmech.2018.06.002

30 Galassi, R. et al. Synthesis and characterization of azolate gold(i) phosphane complexes as thioredoxin reductase inhibiting antitumor agents. Dalton Trans. 41, 5307–5318 (2012). 10.1039/C2DT11781A

31 Galassi, R. et al. A study on the inhibition of dihydrofolate reductase (DHFR) from Escherichia coli by gold (i) phosphane compounds. X-ray crystal structures of (4, 5-dichloro-1 H-imidazolate-1-yl)-triphenylphosphane-gold (i) and (4, 5-dicyano-1 H-imidazolate-1-yl)-triphen. Dalton Transactions 44, 3043–3056 (2015).

32 Molina, D. M. et al. Monitoring Drug Target Engagement in Cells and Tissues Using the Cellular Thermal Shift Assay. Science 341, 84–87 (2013). doi:10.1126/science.1233606

33 Gaetani, M. et al. Proteome Integral Solubility Alteration: A High-Throughput Proteomics Assay for Target Deconvolution. Journal of Proteome Research 18, 4027–4037 (2019). 10.1021/acs.jproteome.9b00500

34 Andor, A. et al. TXNL1 has dual functions as a redox active thioredoxin-like protein as well as an ATP- and redox-independent chaperone. Redox Biology 67, 102897 (2023). 10.1016/j.redox.2023.102897

35 Saei, A. A. et al. Comprehensive chemical proteomics for target deconvolution of the redox active drug auranofin. Redox Biology 32, 101491 (2020). 10.1016/j.redox.2020.101491

36 Sabatier, P. et al. Comprehensive chemical proteomics analyses reveal that the new TRi-1 and TRi-2 compounds are more specific thioredoxin reductase 1 inhibitors than auranofin. Redox Biology 48, 102184 (2021). 10.1016/j.redox.2021.102184

37 Stafford, W. C. et al. Irreversible inhibition of cytosolic thioredoxin reductase 1 as a mechanistic basis for anticancer therapy. Science Translational Medicine 10, eaaf7444 (2018). 10.1126/scitranslmed.aaf7444

38 Gao, J. et al. Structure of the TXNL1-bound proteasome. bioRxiv, 2024.2011.2008.622741 (2024). 10.1101/2024.11.08.622741

39 Ardini, M. et al. The “Doorstop Pocket” In Thioredoxin Reductases─An Unexpected Druggable Regulator of the Catalytic Machinery. Journal of Medicinal Chemistry 67, 15947–15967 (2024). 10.1021/acs.jmedchem.4c00669

40 Takahashi, N. et al. 3D Culture Models with CRISPR Screens Reveal Hyperactive NRF2 as a Prerequisite for Spheroid Formation via Regulation of Proliferation and Ferroptosis. Molecular Cell 80, 828–844.e826 (2020). 10.1016/j.molcel.2020.10.010

41 Ascenzi, F. et al. Identification of a set of genes potentially responsible for resistance to ferroptosis in lung adenocarcinoma cancer stem cells. Cell Death & Disease 15, 303 (2024). 10.1038/s41419-024-06667-w

42 Qiu, B. et al. Phospholipids with two polyunsaturated fatty acyl tails promote ferroptosis. Cell 187, 1177–1190.e1118 (2024). 10.1016/j.cell.2024.01.030

43 Kagan, V. E. et al. Oxidized arachidonic and adrenic PEs navigate cells to ferroptosis. Nature Chemical Biology 13, 81–90 (2017). 10.1038/nchembio.2238

44 Yang, W. S. et al. Peroxidation of polyunsaturated fatty acids by lipoxygenases drives ferroptosis. Proceedings of the National Academy of Sciences 113, E4966–E4975 (2016). 10.1073/pnas.1603244113

45 Yang, Wan S. et al. Regulation of Ferroptotic Cancer Cell Death by GPX4. Cell 156, 317–331 (2014). 10.1016/j.cell.2013.12.010

46 Dierge, E. et al. Peroxidation of n-3 and n-6 polyunsaturated fatty acids in the acidic tumor environment leads to ferroptosis-mediated anticancer effects. Cell Metabolism 33, 1701–1715.e1705 (2021). 10.1016/j.cmet.2021.05.016

47 Liu, X. & Hummon, A. B. Chemical Imaging of Platinum-Based Drugs and their Metabolites. Scientific Reports 6, 38507 (2016). 10.1038/srep38507

48 Friedmann Angeli, J. P., et al. Inactivation of the ferroptosis regulator Gpx4 triggers acute renal failure in mice. Nature cell biology 16, 1180–1191 (2014). 10.1038/ncb3064

49 Davies, S. S. & Guo, L. Lipid peroxidation generates biologically active phospholipids including oxidatively N-modified phospholipids. Chemistry and Physics of Lipids 181, 1–33 (2014). 10.1016/j.chemphyslip.2014.03.002

50 Tyurina, Y. Y. et al. Redox phospholipidomics discovers pro-ferroptotic death signals in A375 melanoma cells in vitro and in vivo. Redox Biology 61, 102650 (2023). 10.1016/j.redox.2023.102650

51 Choi, Y. J. et al. Combined inhibition of IGFR enhances the effects of gefitinib in H1650: a lung cancer cell line with EGFR mutation and primary resistance to EGFR-TK inhibitors. Cancer Chemotherapy and Pharmacology 66, 381–388 (2010). 10.1007/s00280-009-1174-7

52 Moss, D. Y., McCann, C. & Kerr, E. M. Rerouting the drug response: Overcoming metabolic adaptation in KRAS-mutant cancers. Science Signaling 15, eabj3490 10.1126/scisignal.abj3490

53 Humpton, T. J., Hock, A. K., Maddocks, O. D. K. & Vousden, K. H. p53-mediated adaptation to serine starvation is retained by a common tumour-derived mutant. Cancer & Metabolism 6, 18 (2018). 10.1186/s40170-018-0191-6

54 DeNicola, G. M. et al. NRF2 regulates serine biosynthesis in non–small cell lung cancer. Nature Genetics 47, 1475–1481 (2015). 10.1038/ng.3421

55 Fiskus, W. et al. Auranofin Induces Lethal Oxidative and Endoplasmic Reticulum Stress and Exerts Potent Preclinical Activity against Chronic Lymphocytic Leukemia. Cancer Research 74, 2520–2532 (2014). 10.1158/0008-5472.Can-13-2033

56 Torrente, L. et al. Inhibition of TXNRD or SOD1 overcomes NRF2-mediated resistance to β-lapachone. Redox Biology 30, 101440 (2020). 10.1016/j.redox.2020.101440

57 Harris, Isaac S. et al. Glutathione and Thioredoxin Antioxidant Pathways Synergize to Drive Cancer Initiation and Progression. Cancer Cell 27, 211–222 (2015). 10.1016/j.ccell.2014.11.019

58 Campbell, N. K., Fitzgerald, H. K. & Dunne, A. Regulation of inflammation by the antioxidant haem oxygenase 1. Nature Reviews Immunology 21, 411–425 (2021). 10.1038/s41577-020-00491-x

59 DeNicola, G. M. et al. Oncogene-induced Nrf2 transcription promotes ROS detoxification and tumorigenesis. Nature 475, 106–109 (2011). 10.1038/nature10189

60 Weiss-Sadan, T. et al. NRF2 activation induces NADH-reductive stress, providing a metabolic vulnerability in lung cancer. Cell Metabolism 35, 487–503.e487 (2023). 10.1016/j.cmet.2023.01.012

61 Hamamura, R. S. et al. Induction of heme oxygenase-1 by cobalt protoporphyrin enhances the antitumour effect of bortezomib in adult T-cell leukaemia cells. British Journal of Cancer 97, 1099–1105 (2007). 10.1038/sj.bjc.6604003

62 Csongradi, E., doCarmo, J. M., Dubinion, J. H., Vera, T. & Stec, D. E. Chronic HO-1 induction with cobalt protoporphyrin (CoPP) treatment increases oxygen consumption, activity, heat production and lowers body weight in obese melanocortin-4 receptor-deficient mice. International Journal of Obesity 36, 244–253 (2012). 10.1038/ijo.2011.78

63 Kobayashi, S., Harada, Y., Homma, T., Yokoyama, C. & Fujii, J. Characterization of a rat monoclonal antibody raised against ferroptotic cells. Journal of Immunological Methods 489, 112912 (2021). 10.1016/j.jim.2020.112912

64 Zheng, H., Jiang, L., Tsuduki, T., Conrad, M. & Toyokuni, S. Embryonal erythropoiesis and aging exploit ferroptosis. Redox Biology 48, 102175 (2021). 10.1016/j.redox.2021.102175

65 Skoulidis, F. et al. Sotorasib for Lung Cancers with KRAS p.G12C Mutation. New England Journal of Medicine 384, 2371–2381 (2021). doi:10.1056/NEJMoa2103695

66 Wang, X. et al. Identification of MRTX1133, a Noncovalent, Potent, and Selective KRASG12D Inhibitor. Journal of Medicinal Chemistry 65, 3123–3133 (2022). 10.1021/acs.jmedchem.1c01688

67 Holderfield, M. et al. Concurrent inhibition of oncogenic and wild-type RAS-GTP for cancer therapy. Nature 629, 919–926 (2024). 10.1038/s41586-024-07205-6

68 Cregg, J. et al. Discovery of Daraxonrasib (RMC-6236), a Potent and Orally Bioavailable RAS(ON) Multi-selective, Noncovalent Tri-complex Inhibitor for the Treatment of Patients with Multiple RAS-Addicted Cancers. Journal of Medicinal Chemistry 68, 6064–6083 (2025). 10.1021/acs.jmedchem.4c02314

69 Jiang, J. et al. Translational and Therapeutic Evaluation of RAS-GTP Inhibition by RMC-6236 in RAS-Driven Cancers. Cancer Discovery 14, 994–1017 (2024). 10.1158/2159-8290.Cd-24-0027

70 Mohanty, A. et al. Acquired resistance to KRAS G12C small-molecule inhibitors via genetic/nongenetic mechanisms in lung cancer. Science Advances 9, eade3816-eade3816 (2024). 10.1126/sciadv.ade3816

71 Dilly, J. et al. Mechanisms of Resistance to Oncogenic KRAS Inhibition in Pancreatic Cancer. *Cancer Discovery*, OF1-OF27 (2024). 10.1158/2159-8290.CD-24-0177

72 Sattler, M., Mohanty, A., Kulkarni, P. & Salgia, R. Precision oncology provides opportunities for targeting KRAS-inhibitor resistance. Trends in Cancer 9, 42–54 (2023). 10.1016/j.trecan.2022.10.001

73 Luo, J. et al. Overcoming KRAS-Mutant Lung Cancer. American Society of Clinical Oncology Educational Book, 700–710 (2022). 10.1200/EDBK_360354

74 Tanaka, N. et al. Clinical Acquired Resistance to KRASG12C Inhibition through a Novel KRAS Switch-II Pocket Mutation and Polyclonal Alterations Converging on RAS{\textendash}MAPK Reactivation. Cancer Discovery 11, 1913–1922 (2021). 10.1158/2159-8290.CD-21-0365

75 Dunnett-Kane, V., Nicola, P., Blackhall, F. & Lindsay, C. Mechanisms of Resistance to KRAS(G12C) Inhibitors. Cancers 13, 151–151 (2021). 10.3390/cancers13010151

76 Awad, M. M. et al. Acquired Resistance to KRASG12C Inhibition in Cancer. New England Journal of Medicine 384, 2382–2393 (2021). 10.1056/NEJMoa2105281

77 Sang, B. et al. Mechanisms of resistance to active state selective tri-complex RAS inhibitors. bioRxiv, 2025.2004.2024.649345 (2025). 10.1101/2025.04.24.649345

78 Kar, S. et al. Abstract LB281: Mechanisms of resistance to the RAS(ON) multi-selective inhibitor daraxonrasib (RMC-6236) in RAS mutant PDAC and potential resolution with RAS(ON) combination therapies. Cancer Research 85, LB281-LB281 (2025). 10.1158/1538-7445.Am2025-lb281

79 Choe, J. H. et al. Abstract 5507: Mechanisms of resistance to RAS-GTP inhibition in pancreatic cancer. Cancer Research 85, 5507–5507 (2025). 10.1158/1538-7445.Am2025-5507

80 Jiang, L. et al. Ferroptosis as a p53-mediated activity during tumour suppression. Nature 520, 57–62 (2015). 10.1038/nature14344

81 Zhou, Q. et al. Ferroptosis in cancer: from molecular mechanisms to therapeutic strategies. Signal Transduction and Targeted Therapy 9, 55 (2024). 10.1038/s41392-024-01769-5

82 Hangauer, M. J. et al. Drug-tolerant persister cancer cells are vulnerable to GPX4 inhibition. Nature (2017). 10.1038/nature24297

83 Wang, Y. et al. ACSL4 and polyunsaturated lipids support metastatic extravasation and colonization. Cell 188, 412–429.e427 (2025). 10.1016/j.cell.2024.10.047

84 Liang, D. et al. Ferroptosis surveillance independent of GPX4 and differentially regulated by sex hormones. Cell 186, 2748–2764.e2722 (2023). 10.1016/j.cell.2023.05.003

85 Chen, D. et al. iPLA2β-mediated lipid detoxification controls p53-driven ferroptosis independent of GPX4. Nature Communications 12, 3644 (2021). 10.1038/s41467-021-23902-6

86 Dixon, S. J. & Pratt, D. A. Ferroptosis: A flexible constellation of related biochemical mechanisms. Molecular Cell 83, 1030–1042 (2023). 10.1016/j.molcel.2023.03.005

87 Yagoda, N. et al. RAS-RAF-MEK-dependent oxidative cell death involving voltage-dependent anion channels. Nature 447, 865–869 (2007). 10.1038/nature05859

88 Dolma, S., Lessnick, S. L., Hahn, W. C. & Stockwell, B. R. Identification of genotype-selective antitumor agents using synthetic lethal chemical screening in engineered human tumor cells. Cancer Cell 3, 285–296 (2003). 10.1016/S1535-6108(03)00050-3

89 Poursaitidis, I. et al. Oncogene-Selective Sensitivity to Synchronous Cell Death following Modulation of the Amino Acid Nutrient Cystine. Cell Reports 18, 2547–2556 (2017). 10.1016/j.celrep.2017.02.054

90 Anestål, K. & Arnér, E. S. J. Rapid Induction of Cell Death by Selenium-compromised Thioredoxin Reductase 1 but Not by the Fully Active Enzyme Containing Selenocysteine*. Journal of Biological Chemistry 278, 15966–15972 (2003). 10.1074/jbc.M210733200

91 Dixon, S. J. & Stockwell, B. R. The Hallmarks of Ferroptosis. Annual Review of Cancer Biology 3, 35–54 (2019). 10.1146/annurev-cancerbio-030518-055844

92 Hu, K. et al. Suppression of the SLC7A11/glutathione axis causes synthetic lethality in KRAS-mutant lung adenocarcinoma. The Journal of Clinical Investigation 130, 1752–1766 (2020). 10.1172/JCI124049

93 Jiang, H. et al. Ferrous iron–activatable drug conjugate achieves potent MAPK blockade in KRAS-driven tumors. Journal of Experimental Medicine 219 (2022). 10.1084/jem.20210739

94 França, G. S. et al. Cellular adaptation to cancer therapy along a resistance continuum. Nature 631, 876–883 (2024). 10.1038/s41586-024-07690-9

95 Ghandi, M. et al. Next-generation characterization of the Cancer Cell Line Encyclopedia. Nature 569, 503–508 (2019). 10.1038/s41586-019-1186-3

96 Dempster, J. M. et al. Extracting Biological Insights from the Project Achilles Genome-Scale CRISPR Screens in Cancer Cell Lines. bioRxiv, 720243–720243 (2019). 10.1101/720243

97 Dempster, J. M. et al. Chronos: a CRISPR cell population dynamics model. bioRxiv, 2021.2002.2025.432728 (2021). 10.1101/2021.02.25.432728

98 Pacini, C. et al. Integrated cross-study datasets of genetic dependencies in cancer. Nature Communications 12, 1661 (2021). 10.1038/s41467-021-21898-7

99 Babak, M. V. et al. Interfering with Metabolic Profile of Triple-Negative Breast Cancers Using Rationally Designed Metformin Prodrugs. Angewandte Chemie International Edition 60, 13405–13413 (2021). 10.1002/anie.202102266

100 Gazdar, A. F., Girard, L., Lockwood, W. W., Lam, W. L. & Minna, J. D. in Journal of the National Cancer Institute (2010).

101 Chuang, C.-H. et al. Altered Mitochondria Functionality Defines a Metastatic Cell State in Lung Cancer and Creates an Exploitable Vulnerability. Cancer Research 81, 567–579 (2021). 10.1158/0008-5472.Can-20-1865

102. Fellmann, C., et al. An optimized microRNA backbone for effective single-copy RNAi. Cell Reports (2013). 10.1016/j.celrep.2013.11.020

103 Padanad, M. S. et al. Fatty Acid Oxidation Mediated by Acyl-CoA Synthetase Long Chain 3 Is Required for Mutant KRAS Lung Tumorigenesis. Cell Reports 16, 1614–1616 (2016). 10.1016/j.celrep.2016.07.009

104 Abraham, M., Alekseenko, A., Basov, V., Bergh, C., Briand, E., Brown, A., Doijade, M., Fiorin, G., Fleischmann, S., Gorelov, S., Gouaillardet, G., Grey, A., Irrgang, M. E., Jalalypour, F., Jordan, J., Kutzner, C., Lemkul, J. A., Lundborg, M., Merz, P., Lindahl, E. . in Zenodo (2024).

105. Gaussian 16 Rev. C.01 (Wallingford, CT, 2016).

106 Morris, G. M. et al. AutoDock4 and AutoDockTools4: Automated docking with selective receptor flexibility. Journal of Computational Chemistry 30, 2785–2791 (2009). 10.1002/jcc.21256

107 Pettersen, E. F. et al. UCSF Chimera—A visualization system for exploratory research and analysis. Journal of Computational Chemistry 25, 1605–1612 (2004). 10.1002/jcc.20084

108 Salentin, S., Schreiber, S., Haupt, V. J., Adasme, M. F. & Schroeder, M. PLIP: fully automated protein–ligand interaction profiler. Nucleic Acids Research 43, W443–W447 (2015). 10.1093/nar/gkv315

109 Zheng, S. et al. SynergyFinder Plus: Toward Better Interpretation and Annotation of Drug Combination Screening Datasets. Genomics, Proteomics & Bioinformatics 20, 587–596 (2022). 10.1016/j.gpb.2022.01.004

110 Cunniff, B., Snider, G. W., Fredette, N., Hondal, R. J. & Heintz, N. H. A direct and continuous assay for the determination of thioredoxin reductase activity in cell lysates. Analytical Biochemistry 443, 34–40 (2013). 10.1016/j.ab.2013.08.013

111 Ward, N. P. et al. Mitochondrial respiratory function is preserved under cysteine starvation via glutathione catabolism in NSCLC. Nature Communications 15, 4244 (2024). 10.1038/s41467-024-48695-2

112 Cox, Andrew G. et al. Mitochondrial peroxiredoxin 3 is more resilient to hyperoxidation than cytoplasmic peroxiredoxins. Biochemical Journal 421, 51–58 (2009). 10.1042/bj20090242

113 Gaetani, M. & Zubarev, R. A. in Cell-Wide Identification of Metabolite-Protein Interactions (eds Aleksandra Skirycz, Marcin Luzarowski, & Jennifer C. Ewald) 91–106 (Springer US, 2023).

114 Zhang, X., Lytovchenko, O., Lundström, S. L., Zubarev, R. A. & Gaetani, M. Proteome Integral Solubility Alteration (PISA) Assay in Mammalian Cells for Deep, High-Confidence, and High-Throughput Target Deconvolution. Bio-protocol 12, e4556 (2022). 10.21769/BioProtoc.4556

115 Dobin, A. et al. STAR: ultrafast universal RNA-seq aligner. Bioinformatics 29, 15–21 (2012). 10.1093/bioinformatics/bts635

116 Liao, Y., Smyth, G. K. & Shi, W. FeatureCounts: An efficient general purpose program for assigning sequence reads to genomic features. Bioinformatics (2014). 10.1093/bioinformatics/btt656

117 Ewels, P., Magnusson, M., Lundin, S. & Käller, M. MultiQC: summarize analysis results for multiple tools and samples in a single report. Bioinformatics 32, 3047–3048 (2016). 10.1093/bioinformatics/btw354

118 Xie, Z. et al. Gene Set Knowledge Discovery with Enrichr. Current Protocols 1, e90–e90 (2021). 10.1002/cpz1.90

119 Kuleshov, M. V. et al. Enrichr: a comprehensive gene set enrichment analysis web server 2016 update. Nucleic acids research (2016). 10.1093/nar/gkw377

120 Chen, E. Y. et al. Enrichr: Interactive and collaborative HTML5 gene list enrichment analysis tool. BMC Bioinformatics (2013). 10.1186/1471-2105-14-128

121 Kang, Y. P. & DeNicola, G. M. in Metabolic Reprogramming: Methods and Protocols (eds Salvatore Papa & Concetta Bubici) 51–63 (Springer US, 2023).

122 Kang, Y. P. et al. Non-canonical Glutamate-Cysteine Ligase Activity Protects against Ferroptosis. Cell Metabolism 33, 174–189.e177 (2021). 10.1016/j.cmet.2020.12.007

