## Supplementary Information for "Thioredoxin Reductase 1 inhibition triggers ferroptosis in KRAS-independent lung cancers"

**Supplementary figure legends**

**Supplementary Figure 1. Knockdown of TRXR1 is lethal for KRAS-WT and EGFR-MUT LC.** A) Crystal violet assay and (B) immunoblot of cells transfected with the indicated siRNAs (72 hours post transfection. C) Representative images of LIVE/DEAD assay in the indicated 3D spheroids (n=3 independent replicates) and (D) quantification performed on the indicated cell lines. +Dox, indicates doxycycline to activate the shRNA; CTRL, control. P values were calculated using one-way ANOVA followed by Sidak's tests. E) Immunoblot for TRXR1 in the indicated spheroids with or without shRNA induction (representative of n=2 independent shRNAs).

**Supplementary Figure 2. The gold(I) compounds are reversible allosteric inhibitors of TRXR1.** A) Docking of CS47 and DM20 cluster 2 with the human TRXR1 (PDB. 2CFY and 1W1C). The protein is represented in dimeric quaternary structure; a molecular surface is represented to better highlight the allosteric binding cleft. NADPH and FAD are represented as green and orange VdW spheres. Gold as VdW yellow sphere. Closeup at the bottom shows the interactions with residues (Thr286, Pro190, Ile178, Ser185, Leu186, Leu183, Thr263, Asp182) in this site that are mostly not specific (no ion-ion or H-bonding) and the site is on the surface, not deeply inserted into the protein surface. B) Immunoblot and (C) quantification of CETSA assay in H522 cells showing stabilization of TRXR1 in CS47 condition. P values represent multiple t test as a given temperature (n=2 independent experiments). D) PISA assay performed in H522 cells treated as indicated. Red and blue dots indicate significant hits in both conditions. E) Venn diagrams showing the overlap between the significant PISA hits at 5 μM CS47 condition of those identified by Sabatier et al. PMID: 34788728 for auranofin and TRi-1 and TRi-2.

**Supplementary Figure 3. $^{1}$H NMR spectra of the gold(I) compounds.** A) $^{1}$H NMR spectra and chemical shifts of CS47 and (B) DM20 over 72 hours. Ppm, part per million.

**Supplementary Figure 4. ESI-MS spectra of the gold(I) compounds.** A) ESI-MS (+) spectra and ion annotations of CS47 and (B) DM20 freshly prepared and at 24 hours in methanol solution (C, D). The m/z range is 100-1800.

**Supplementary Figure 5. TRXR1 activity, cell viability, and redox status of TRX1 and PRDX1 in KRAS-WT and KM lung cancer cells.** A) Assay schematic and quantification of the TRXR1 activity measured via a DNTP-based kit and (B) via consumption of NADPH. P values were measure via two-way ANOVA followed by Dunnett's multiple comparisons with * P<0.05, ** P<0.01, *** P<0.001, **** P<0.0001. C) MTT viability curves of the indicated LC cell lines treated for 48 hours as reported. D) Schematics of the TRXR/TRX1/PRDX1 redox pathways and redox immunoblot showing the reduced and oxidized forms of TRX1 and PRDX1 upon TRXR1 inhibition in the indicated cell lysates (n=2 independent experiments). Notice that oxidation and oligomer formation happen only in sensitive KRAS-WT cells. HMW, high molecular weight.

**Supplementary Figure 6. TRXR1 inhibition induces ferroptosis in KRAS-WT LC cells.** A) Malondialdehyde (MDA) quantification in the indicated cell lines. Data are represented as fold change over vehicle for each cell line (n= 4 replicates). P values indicate unpaired two-tailed student t test. B) Representative pictures and quantification of $Fe^{2+}$ using FerroOrange in H522 cells. P values indicate unpaired two-tailed student t test (n=3 replicates). C) Synergy plots of Fer-1 or DFO with TRXR1 inhibition in H661. (D) Synergy plots of L-cysteine or NAC with TRXR1 inhibition of the indicated KRAS-WT LC lines at 48 hours (n=2 independent experiments/each combination). P values for the average synergy scores were derived by bootstrapping of the dose–response matrix in www.synergyfinderplus.org.

**Supplementary Figure 7. CS47 detection, pharmacokinetics and toxicity *in vivo*.** A) Schematic of the derivatization method and detection of CS47 in plasma of NCG mice. B) Pharmacokinetic (PK) curve of CS47 in mouse plasma and C) PK parameters. Terminal elimination T1/2, terminal elimination half-life; $T_{max}$, time to reach maximum concentration; $C_{max}$, maximum concentration; $AUC_{last}$, area under the concentration-time curve from 0 until the last quantifiable concentration; $V_z/F$, apparent volume of distribution during the terminal phase after non-intravenous administration; CL/F, apparent total body clearance of drug from plasma after oral administration; MRT, mean residence time. D) Assessment of body weight, (E) white blood cells (WBC), (F) platelets and erythrocytes of mice bearing H522 xenografts treated as indicated. G) H&E staining of lung, liver and kidneys of mice bearing H522 xenografts treated as indicated. H) Pathological lung examination of mice bearing H522 xenografts treated as indicated. P values indicate two-way ANOVA followed by Dunnett's multiple tests.

**Supplementary Figure 8. TRXR1 inhibition in mice bearing EGFR-MUT LC xenografts.** A) Tumor growth curves, schematics and post-dissection images of H1650 xenografts (n=5-6 mice/group) treated as indicated. P values represent two-way ANOVA followed by Dunnett's multiple comparisons with ns P>0.05, **** P<0.0001. B) Assessment of body weight and (C) WBC of mice bearing H1650 xenografts. P values indicate two-way ANOVA followed by Dunnett's multiple comparisons.

**Supplementary Figure 9. TRXR1 inhibition transcriptionally reprograms lipid, GSH and iron metabolism.** A) Gene expression of the indicated genes in KM (H157, H1792, H460, A549), KRAS-WT (H522, H661, H1299, H1993) and EGFR-MUT (PC9, H1650, HCC827) LC cell lines treated as indicated for 24 hours. B) Volcano plot showing the expression proteomics hits and C) corresponding significant

Enrichr pathways of H522 cells treated with CS47 and compared to vehicle. Legend represents $-Log_{10}$ (P value)

**Supplementary Figure 10. GSH synthesis and supplementation compensate for TRXR1 inhibition.** A) Schematic of glutathione synthesis and steady-state metabolomics of the metabolites involved in GSH synthesis in H522 and A549 cell lines treated as indicated. P values indicate unpaired two-tailed student t test *P<0.05, **P<0.01, ***P<0.001, ****P<0.0001. B) Quantification of GSH, GSSG, and GSH/GSSG in the indicated cell lines after 24 hours of treatments. P values were calculated using unpaired two-tailed student t test. C) Synergy plots of TRXR1 inhibition with GSH-EE in KRAS-WT LC cells and (D) with BSO in the indicated KM LC lines at 48 hours (n=2 independent experiments/each combination). P values for the average synergy scores were derived by bootstrapping of the dose–response matrix in www.synergyfinderplus.org.

**Supplementary Figure 11. Direct NRF2 activation alone is not sufficient to induce cell death and HMOX1.** A) Synergy plots of TBHQ and KI696 with TRXR1 inhibition in KRAS-WT LC at 48 hours (n=2 independent experiments/each combination). P values for the average synergy scores were derived by bootstrapping of the dose–response matrix in www.synergyfinderplus.org B) Pearson correlation the DepMap genetic dependencies on TXNRD1 and NFE2L2 or (C) KEAP1 (DepMap Public 23Q4+Score, Chronos) in LC cell lines. D) Immunoblot for NRF2 targets in the lysates of the indicated cell lines, treated as indicated for 48 hours.

**Supplementary Figure 12. HMOX1 is required to execute ferroptosis downstream TRXR1 inhibition.** A) Immunoblot of H661 cells stably transfected and sorted for high (hi) and low expression (lo) of HMOX1-GFP (n=2 independent experiments). B) MTT viability assay and IC50 values of the indicated cell lines treated with CS47 and auranofin for 48 hours. C) Synergy plots of Cobaltic Protoporphyrin IX Chloride (CoPP) with TRXR1 inhibition in KRAS-WT LC cells at 48 hours (n=2 independent experiments/each combination). P values for the average synergy scores were derived by bootstrapping of the dose–response matrix in www.synergyfinderplus.org. D) Representative images and E) quantification of $Fe^{2+}$ using FerroOrange. F) Representative images and G) quantification of number of cells treated with CS47 alone or in combination with *shHMOX1* (n=2 independent experiments). P values were calculated using unpaired two-tailed student t test. (H) Immunoblot for HO-1 in the indicated cell lines treated with CS47 alone or in combination with *shHMOX1* (n=2 independent experiments).

**Supplementary Figure 13. ERK and HO-1 activation in parental and RMCR cells.** A) Immunoblot in parental and (B) RMCR cells treated for 4 days as indicated.

**Supplementary Figure 1**

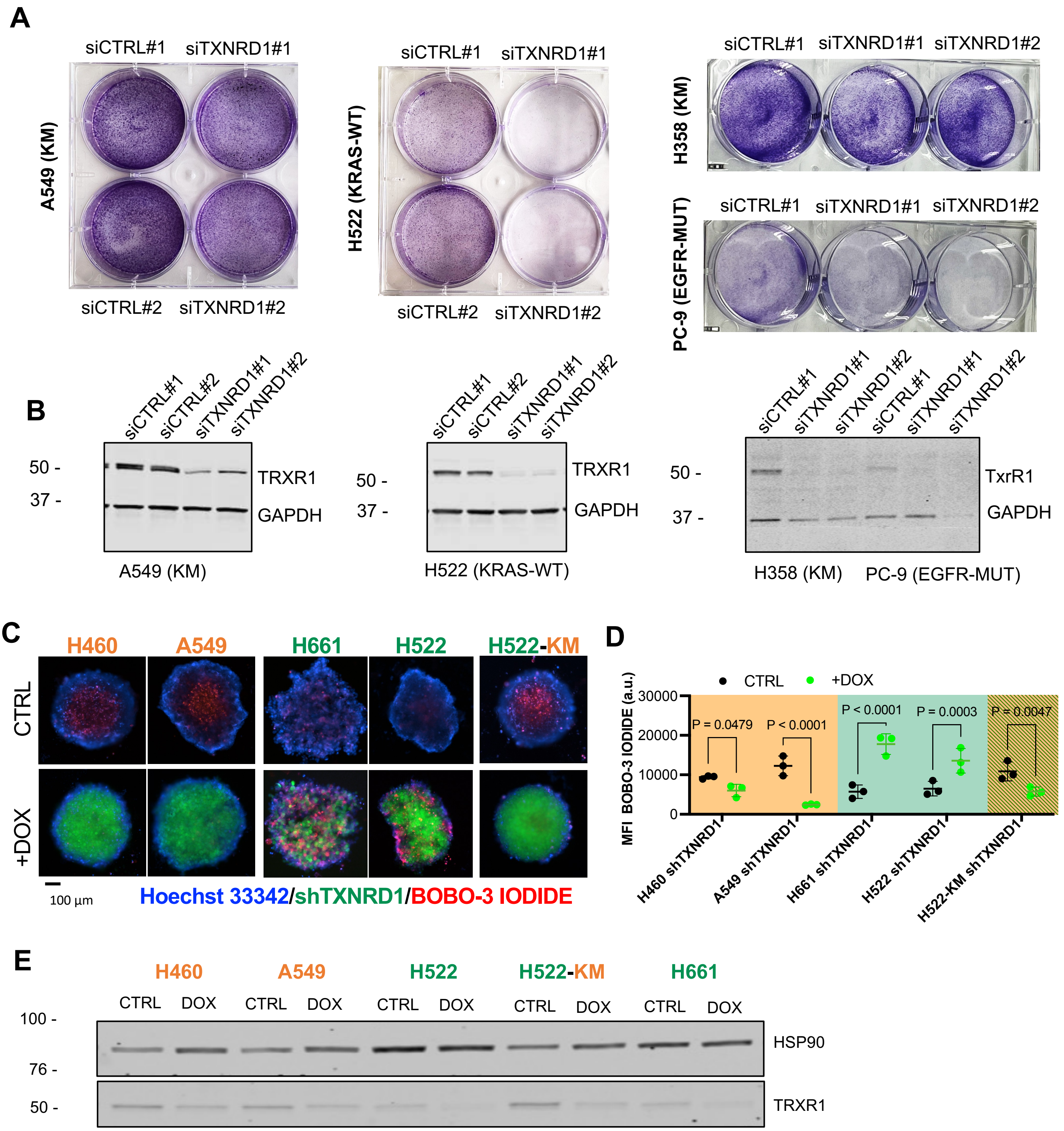

### Supplementary Figure 2

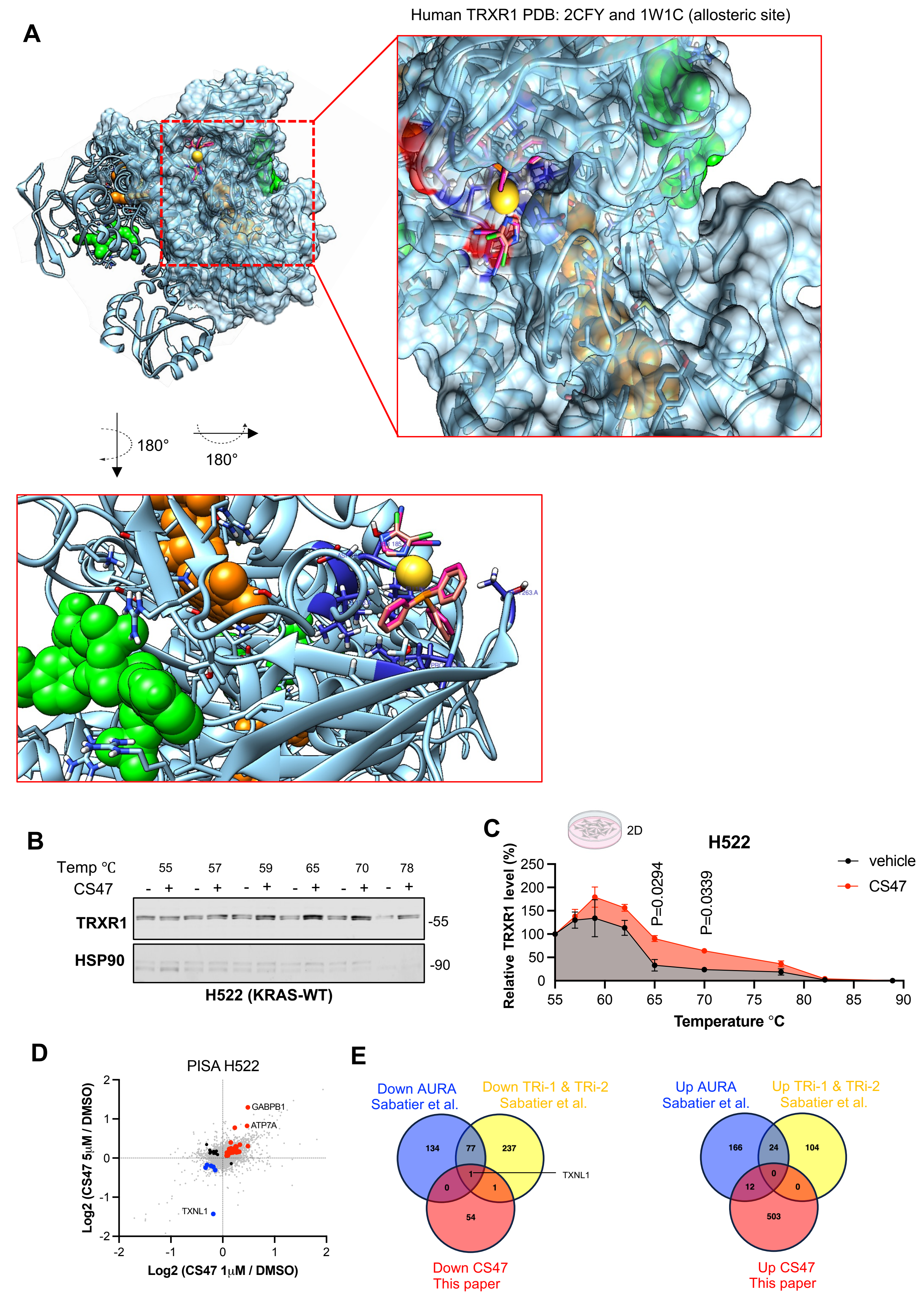

### Supplementary Figure 3

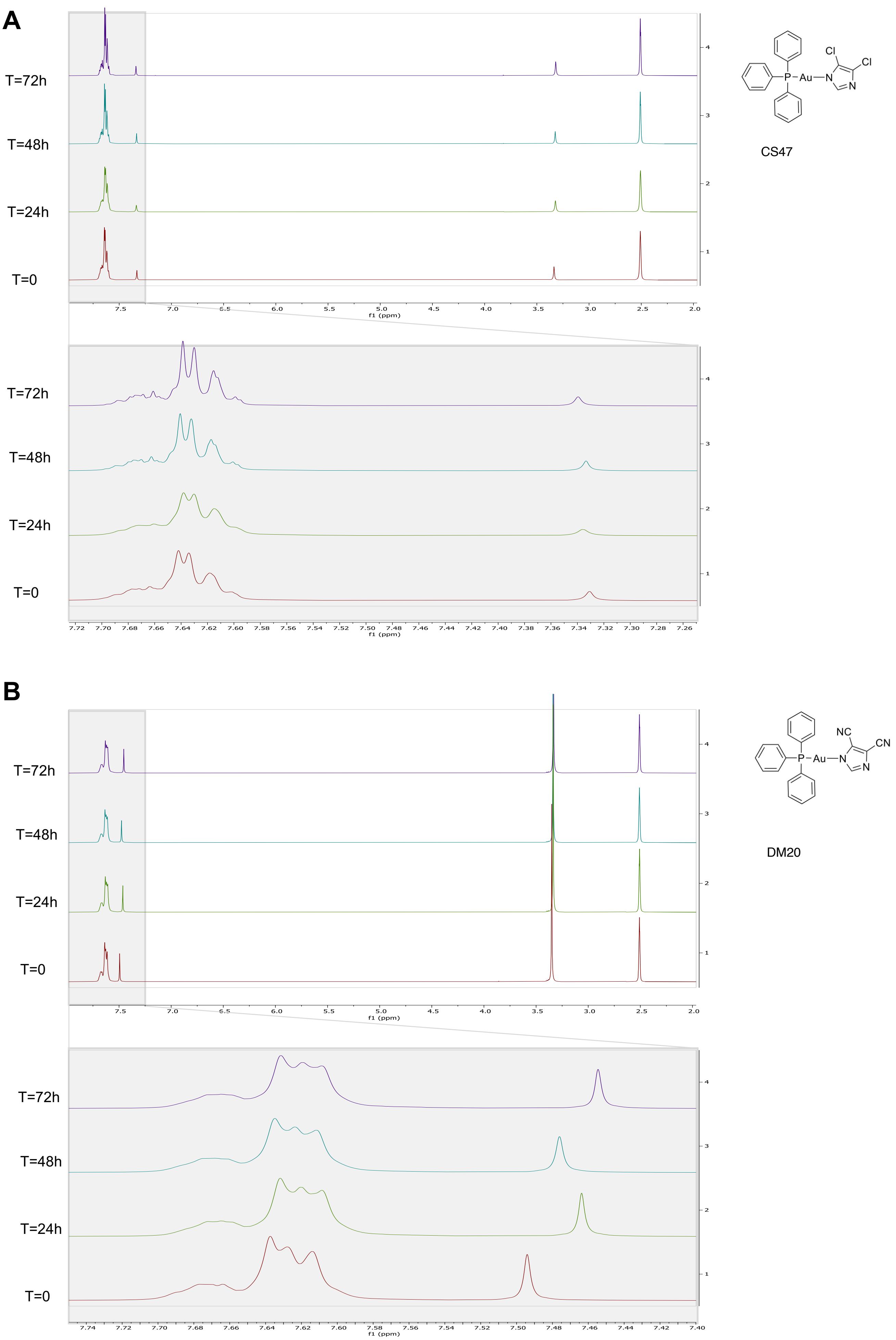

### Supplementary Figure 4

**A**

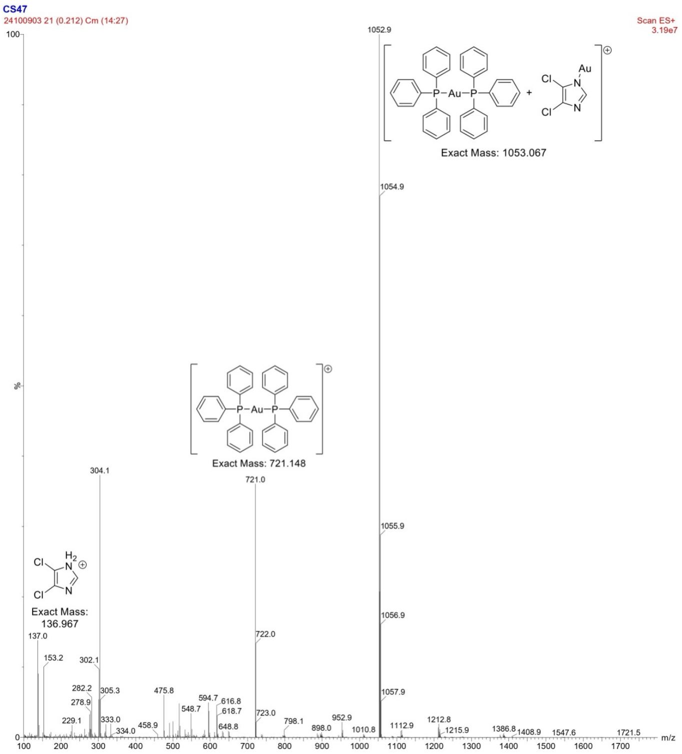


**B**

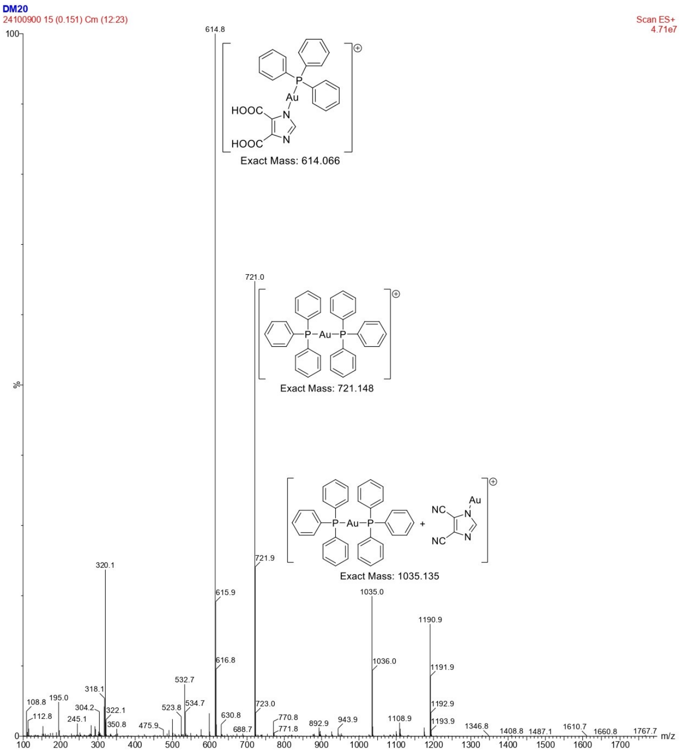


**C**

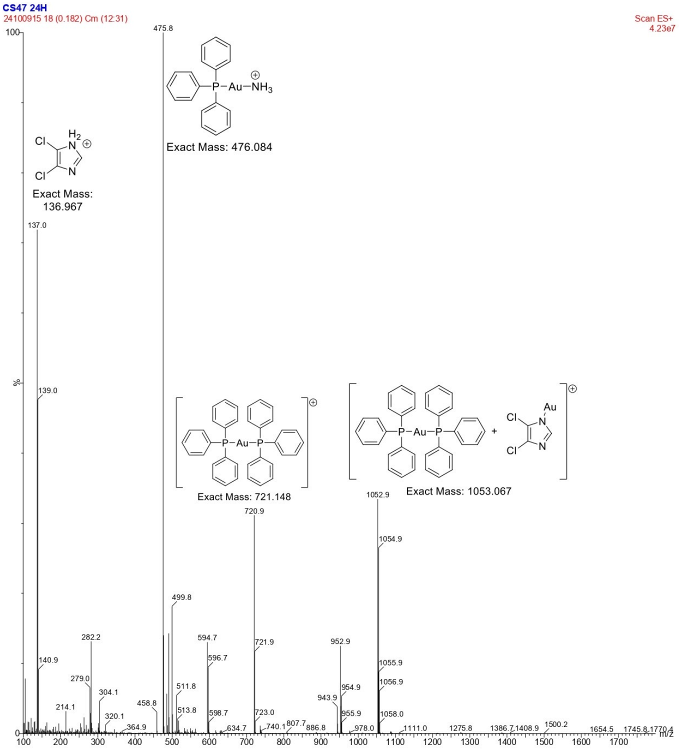


**D**

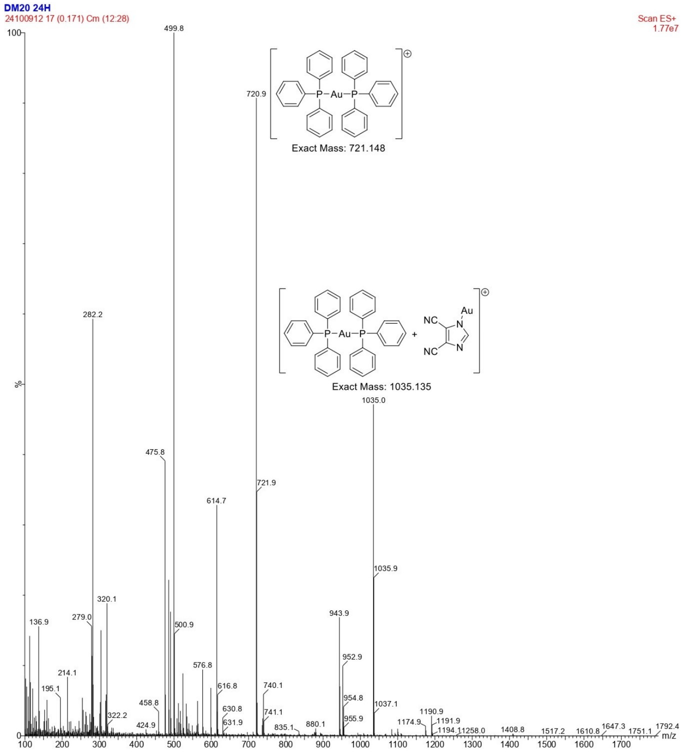

#### Supplementary Figure 5

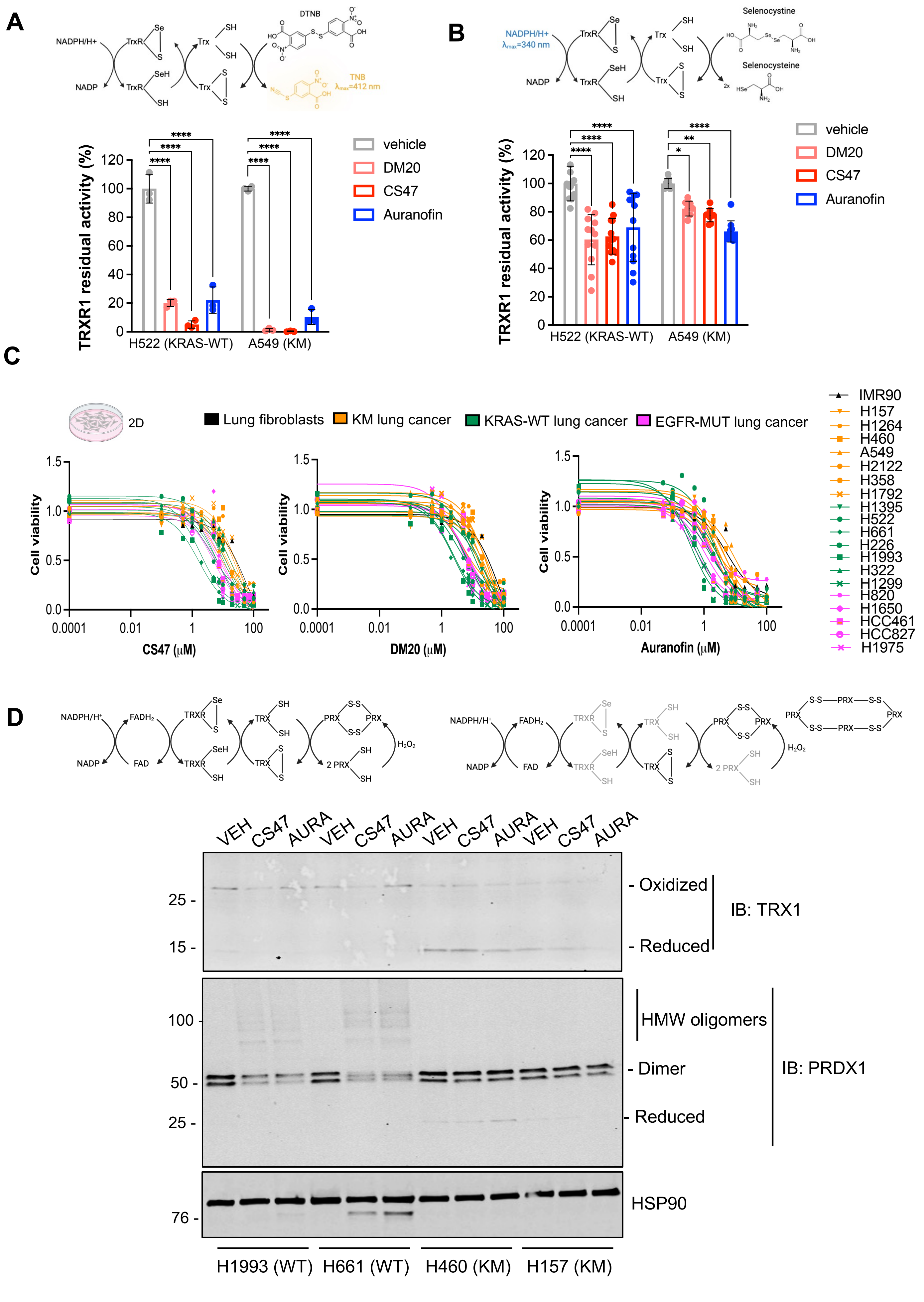

### Supplementary Figure 6

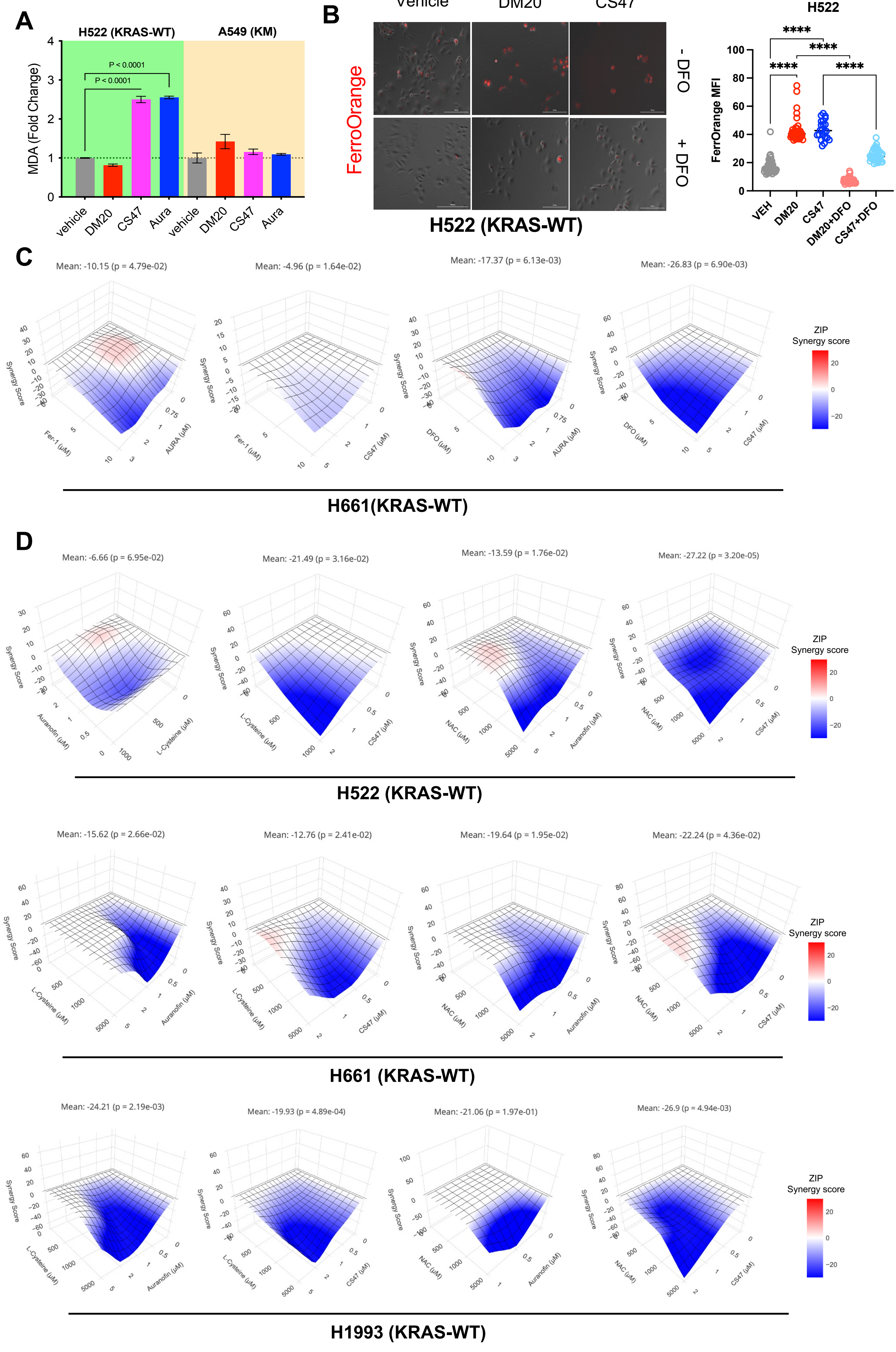

### Supplementary Figure 7

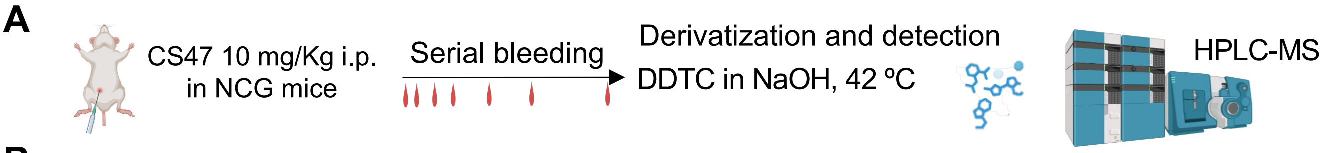


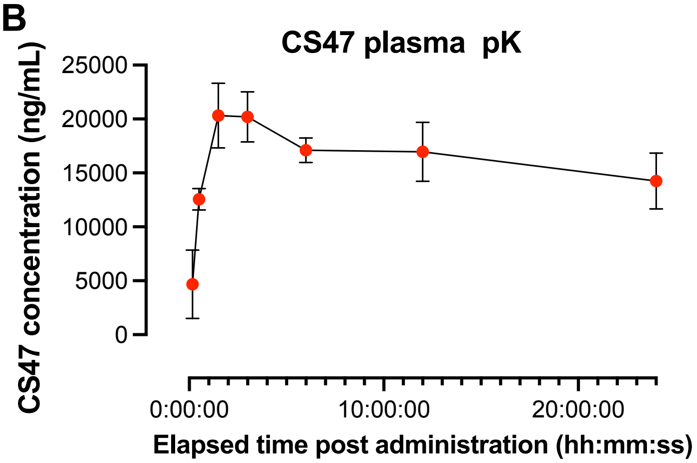


C

| PK Parameters | |
|---|---|
| Terminal T½ (min): | 3803 |
| $T_{max}$ (min): | 90 |
| $C_{max}$ (ng/mL): | 20333 |
| $AUC_{last}$ (min*ng/mL): | 23,737,567 ± 989,987 |
| $V_z/F$ (mL/kg): | 538 |
| CL/F (mL/min/kg): | 0.098 |
| MRT (min): | 686 |

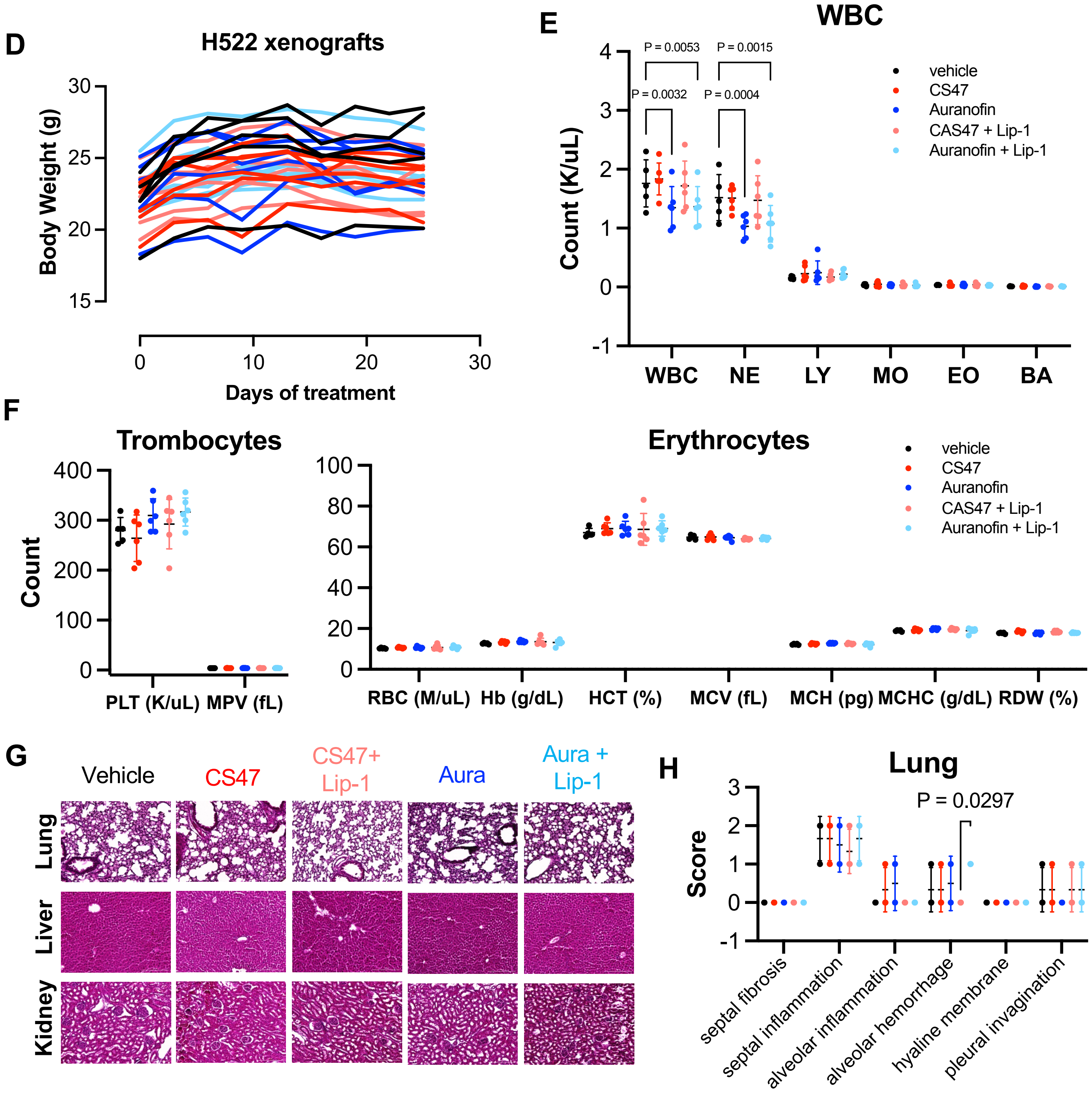

#### Supplementary Figure 8

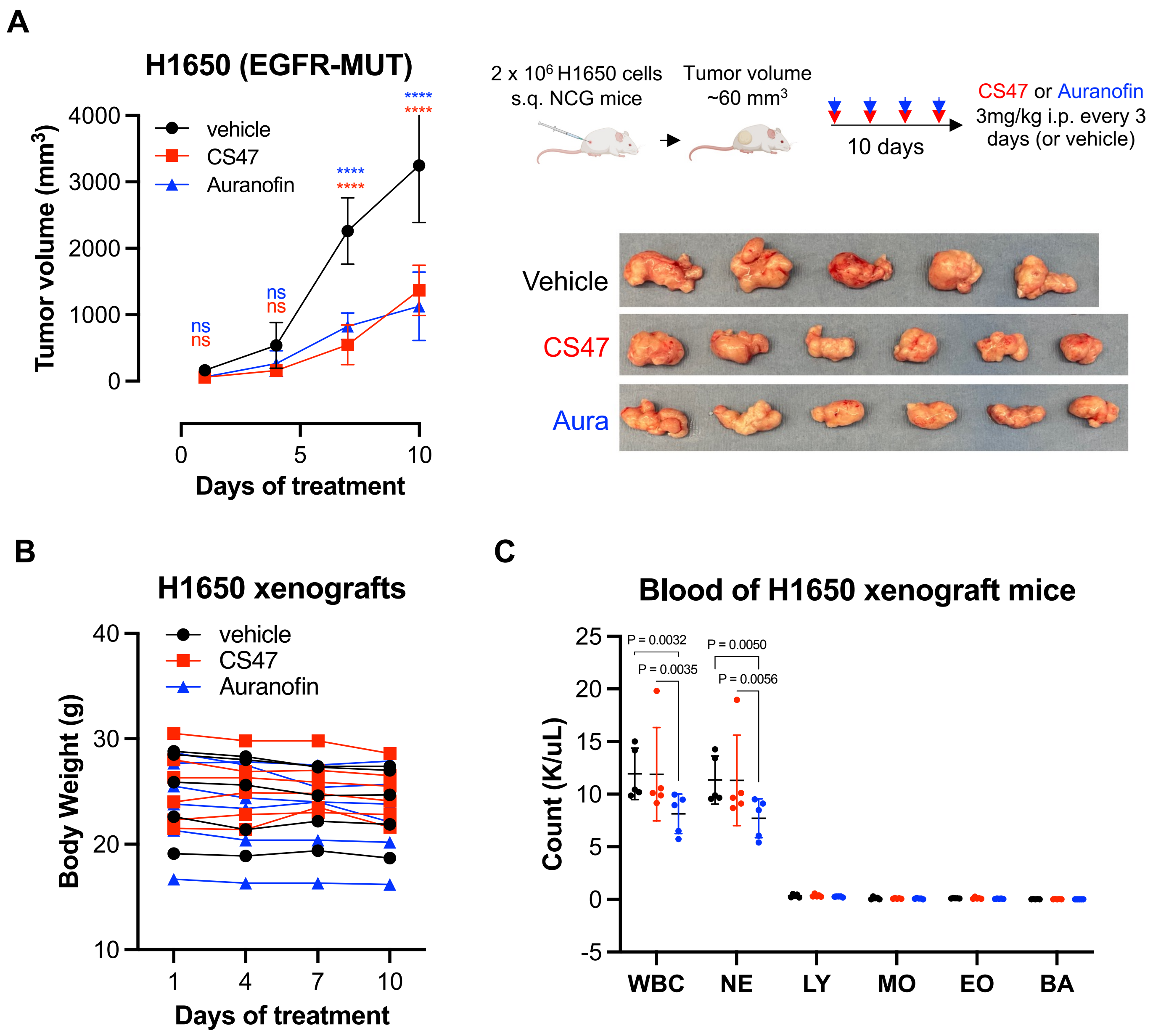

#### Supplementary Figure 9

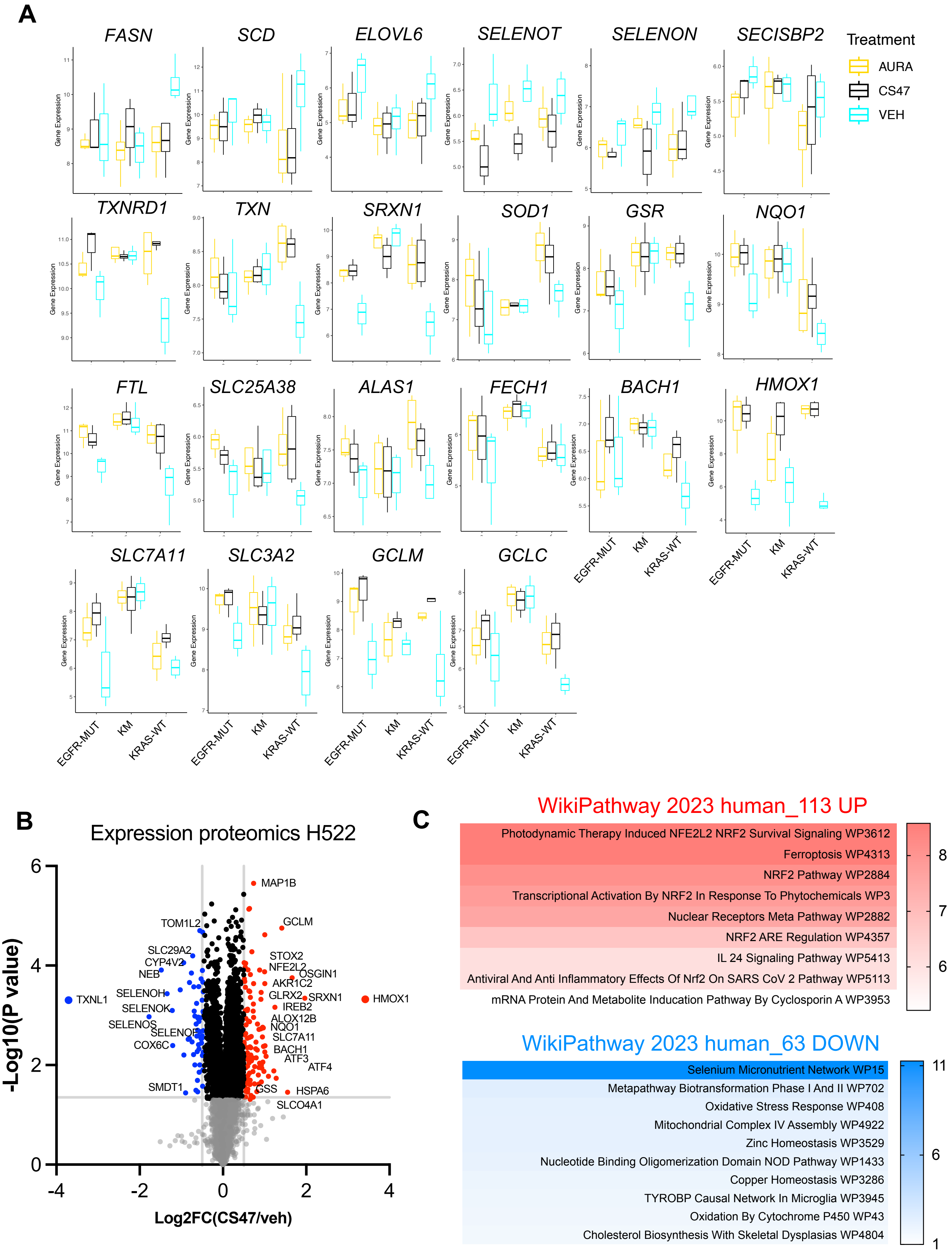

### Supplementary Figure 10

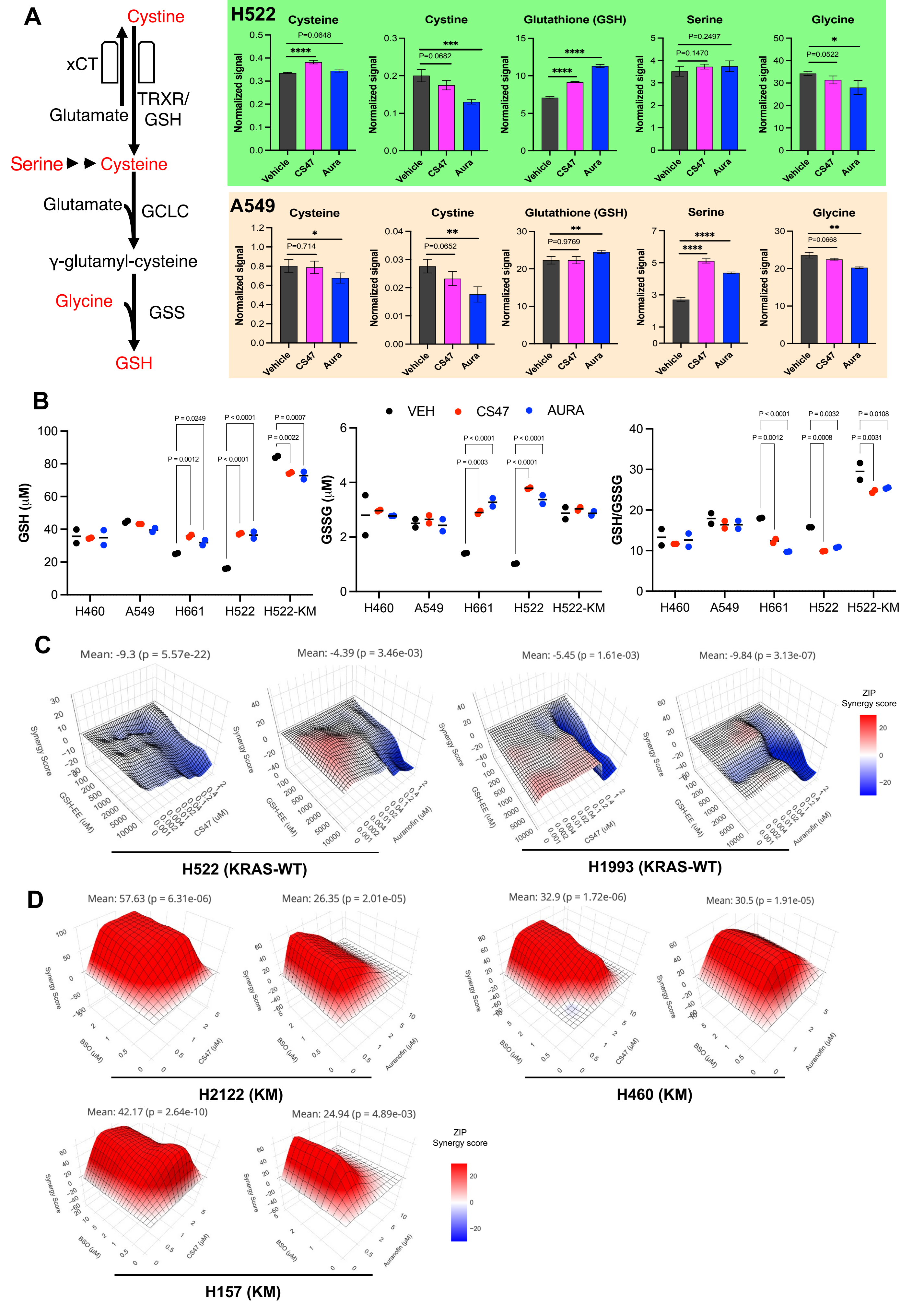

### Supplementary Figure 11

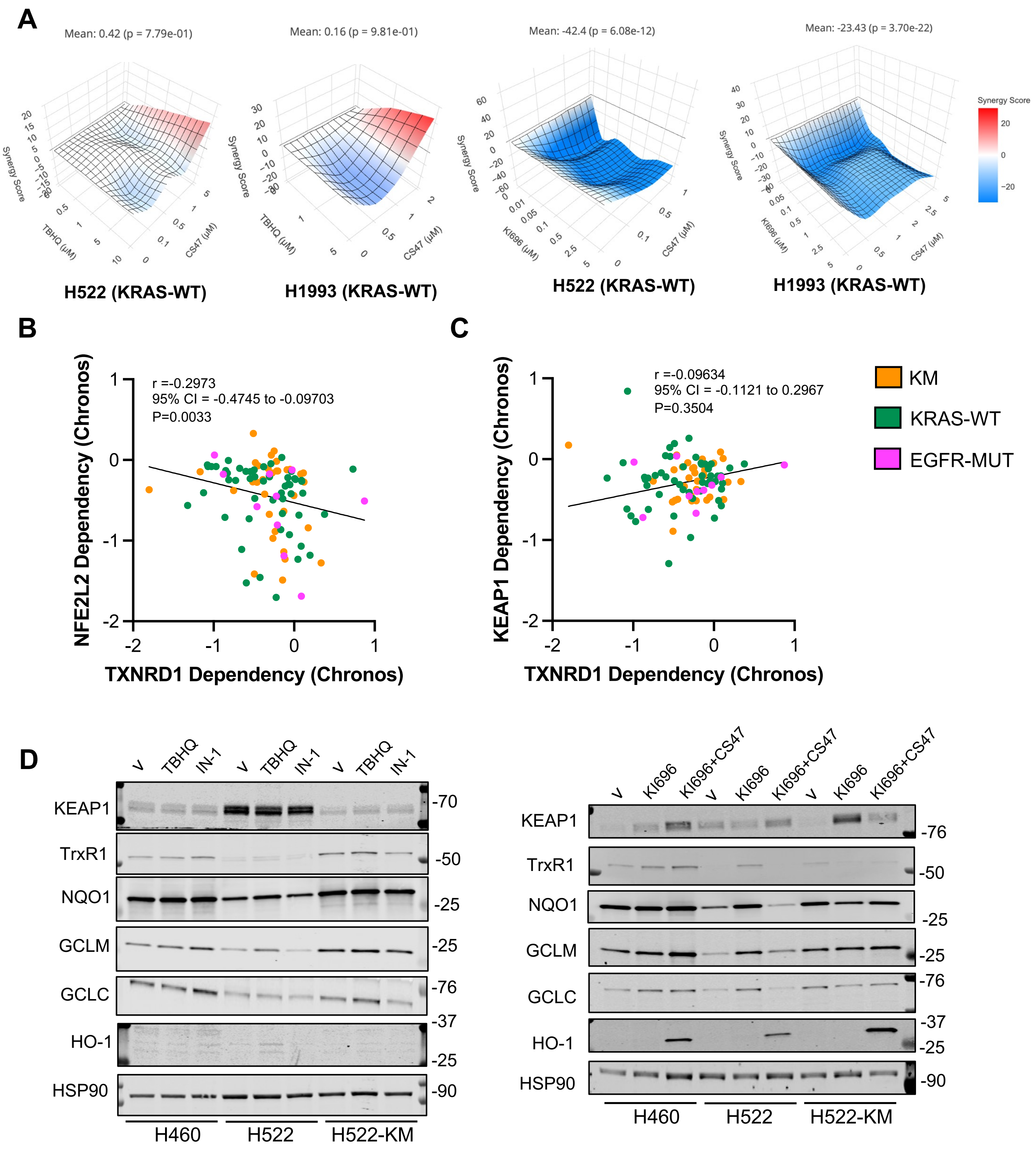

### Supplementary Figure 12

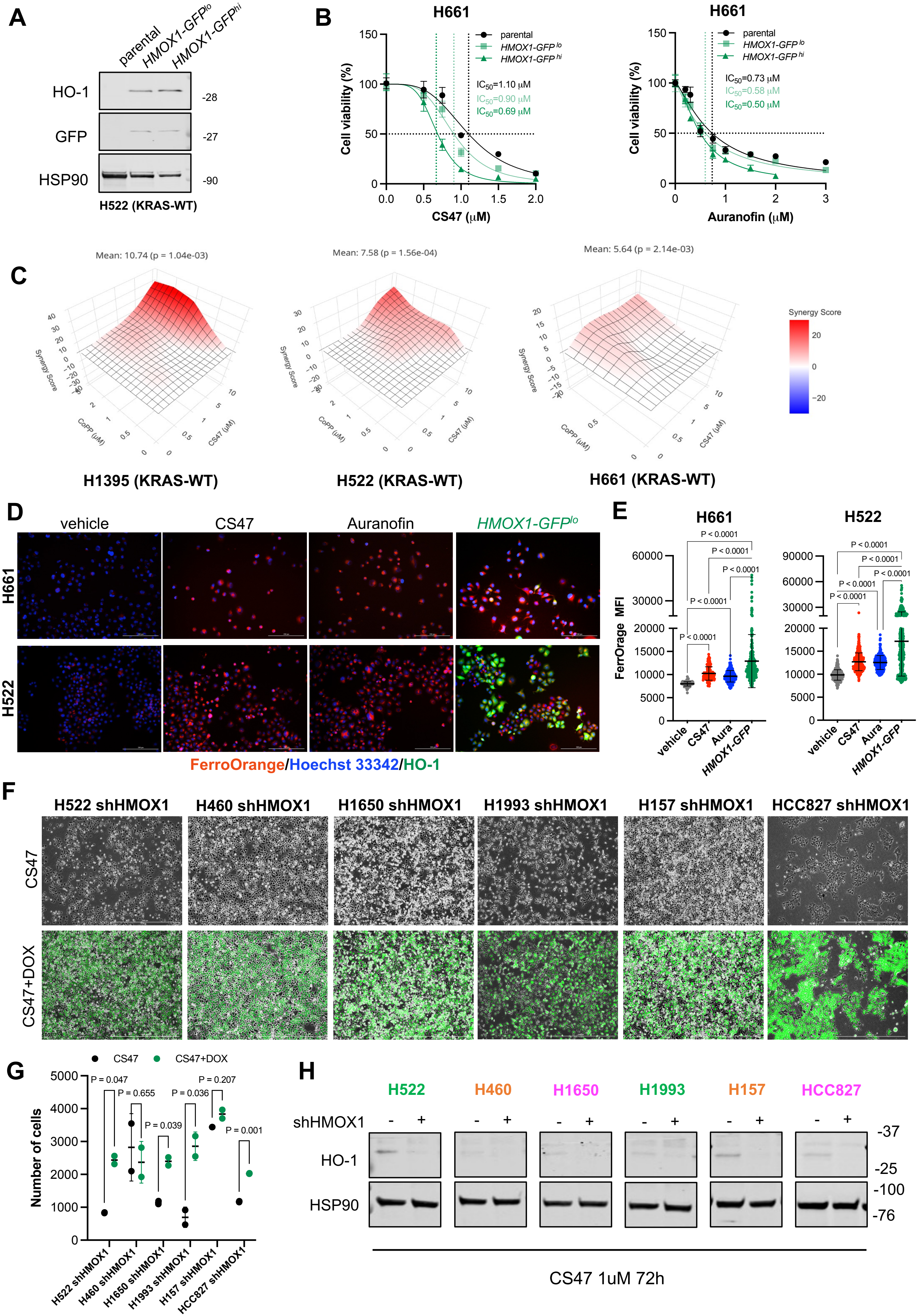

**Supplementary Figure 13**

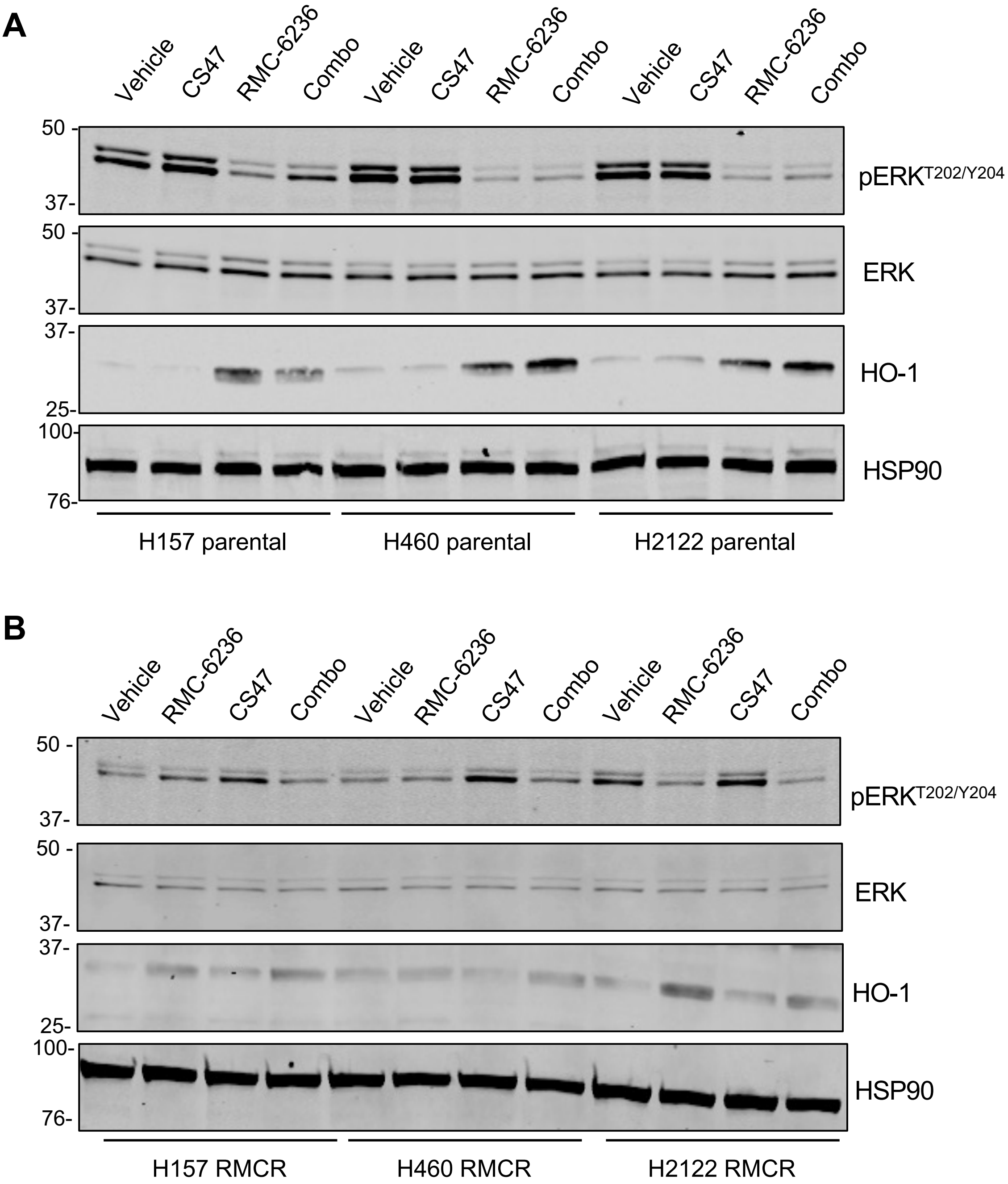

#### Supplementary Table 1. CS47 Acute toxicity experiment

| | Mouse ID | BW (g) | Acute Observation | 30 min Observation | 8 hours Observation | 24 hours Observation | BW (g) at 24 hours | 25 days Observation |
|---|---|---|---|---|---|---|---|---|
| **10 mg/kg (1 mg/ml, 10 ml/kg)** | 2-2 | 23.5 | No ill effects | Reduced activity | Slightly reduced activity | Normal activity | 22.1 | Deceased |
| | 2-3 | 22.5 | No ill effects | Reduced activity | Slightly reduced activity | Normal activity | 22.2 | Deceased |
| **5 mg/kg (0.5 mg/ml, 10 ml/kg)** | 1-3 | 21.2 | No ill effects | Reduced activity | Normal activity | Normal activity | 22.2 | Normal activity |
| | 1-1 | 22.8 | No ill effects | Normal activity | Normal activity | Normal activity | 23.1 | Normal activity |
| **3 mg/kg (0.3 mg/ml, 10 ml/kg)** | 1-a1 | 22.6 | No ill effects | Normal activity | Normal activity | Normal activity | 22.8 | Normal activity |
| | 1-a2 | 22.7 | No ill effects | Normal activity | Normal activity | Normal activity | 23.2 | Normal activity |
| **Vehicle** | 3-1 | 21.2 | No ill effects | - | - | Normal activity | 21.5 | Normal activity |
| | 3-2 | 22.0 | No ill effects | - | - | Normal activity | 22.8 | Normal activity |
| | 3-3 | 21.0 | No ill effects | - | - | Normal activity | 22.0 | Normal activity |
